# Mutational constraints on RSV F and its neutralization by antibodies

**DOI:** 10.64898/2026.02.12.705519

**Authors:** Cassandra A.L. Simonich, Teagan E. McMahon, Gavin Juviler, Lucas Kampman, Helen Y. Chu, Jesse D. Bloom

## Abstract

New antibodies targeting the F protein of RSV have substantially reduced infant hospitalizations^1^. However, viral resistance is a concern: one antibody failed clinical trials due to a resistant strain^2^, and sporadic resistance mutations to the most widely used antibody (nirsevimab) have been identified^3–6^. Here we define how RSV F mutations affect antibody neutralization. We first provide a biophysical model of how the buffering of bivalent IgG binding combines with the lower Fab potency of nirsevimab to subtype B to make resistance to this antibody more common in subtype B than A strains. We then perform pseudovirus deep mutational scanning to safely measure how nearly all mutations to F affect its cell entry function and neutralization by IgG and Fab forms of nirsevimab, clesrovimab, and several other key antibodies. We use these measurements to enable real-time surveillance of RSV sequences for antibody resistance, and show that resistant strains have arisen sporadically but are currently rare. Overall, our work shows how Fab potency and epitope specificity combine to determine how viral mutations impact antibody neutralization, enables monitoring for natural RSV strains resistant to antibodies of public-health importance, and can help guide development of future antibodies with resilience to viral escape.

## Introduction

Respiratory syncytial virus (RSV) is a leading cause of infant hospitalization in the United States and infant mortality globally^7,8^. Two monoclonal antibodies, nirsevimab and clesrovimab, were recently licensed and recommended to prevent severe RSV disease in infants^9^. Nirsevimab has demonstrated ~75-80% effectiveness against RSV-associated hospitalizations of infants^10^ and (along with maternal vaccination) has reduced infant hospitalizations in countries where it is widely used^1^. Additional antibodies are under development including for low- and middle-income countries where RSV is a major cause of infant mortality^11,12^.

Nirsevimab and clesrovimab target the prefusion conformation of the RSV fusion protein (F)^13,14^, binding the apex antigenic region Ø and the lateral face antigenic region IV, respectively. While F is less variable than the surface proteins of influenza and SARS-CoV-2, it still evolves sufficiently fast to pose challenges for monoclonal antibodies^15,16^. Regeneron’s anti-F antibody suptavumab failed a Phase 3 clinical trial from 2015-2017 because it was escaped by mutations in circulating RSV-B strains^2^. While nirsevimab has been in use in the United States for three years and retained high effectiveness, sporadic resistant strains have been identified, particularly in breakthrough infections^3–6,16–20^. Therefore, surveillance of RSV sequences for potential escape mutations is an important public health priority^4,6,20,21^. However, surveillance is hindered by incomplete knowledge of how F mutations affect key antibodies, as prior studies have identified only a limited set of resistance mutations by passaging lab-adapted RSV in the presence of antibody^5,14,22^ or looking for mutations in antibody binding footprints^4–6,17,20^.

Here we provide a complete quantitative understanding of how RSV mutations affect antibodies by combining a biophysical model of bivalent IgG neutralization with deep mutational scanning measurements of how all amino-acid mutations affect F’s cell entry function and neutralization by the IgG and Fab forms of key antibodies. Our work explains why nirsevimab resistance mutations are more common in one of the two RSV subtypes, defines the potential for escape from existing and candidate clinical antibodies, and informs a sequence-surveillance platform to rapidly identify antibody-resistant natural RSV strains.

## Results

### Bivalent IgG binding buffers resistance

Surveillance of individuals who received prophylaxis and experimental studies performed for resistance analysis for FDA licensure have found that viral mutations that escape nirsevimab are more common and impactful in subtype B than subtype A RSV strains^3–6,16–20^. This difference has been puzzling, since nirsevimab has a near-identical binding footprint on the F proteins of subtypes A and B^13^, which have similar structures^23,24^ and ~90% protein sequence identity^25^.

We hypothesized that the greater prevalence of escape mutations in RSV B is because nirsevimab binds with lower affinity to the F from subtype B versus A^19,26^. IgG antibodies, which are produced by the human body and are the form in which nirsevimab is administered, have two antigen-binding arms, but antibodies can be artificially produced in a Fab form with just one binding arm. We built a biophysical model of how mutations that reduce the Fab affinity for a viral antigen impact neutralization by both Fab and bivalent IgG (Supplementary Appendix). The key insight is that when the Fab affinity is sufficiently high, viral mutations that decrease Fab affinity do not measurably decrease IgG neutralization because bivalent binding keeps nearly all the IgG bound to even the mutant virus^27–29^ (Figure 1a). We used the model to predict how the IgG and Fab forms of an antibody would neutralize a virus to which the Fab had high or moderate affinities, using parameter estimates that roughly reflect nirsevimab against RSV subtypes A and B (Figure 1b).

**Figure 1.**
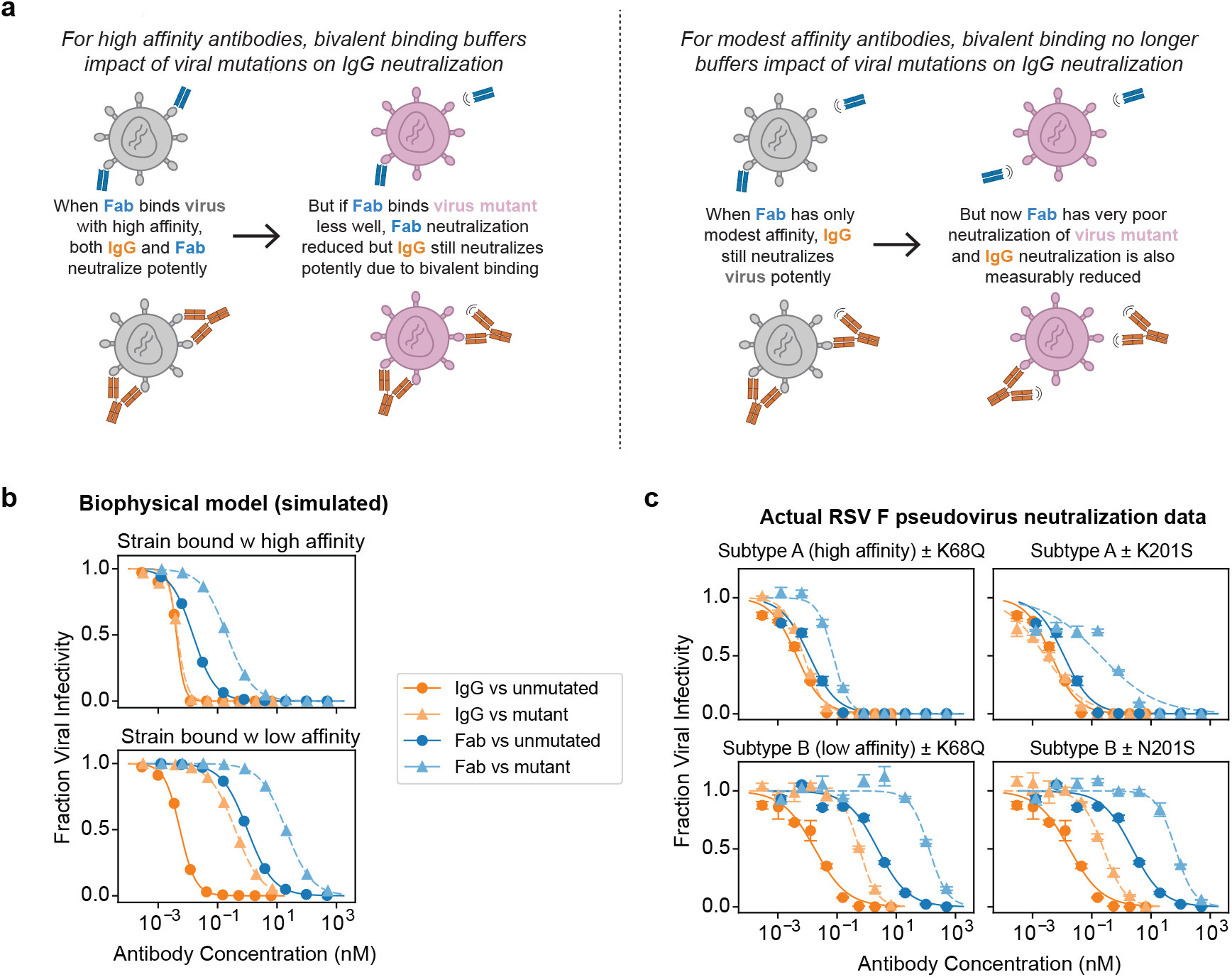
RSV F mutations cause more nirsevimab escape in subtype B versus A because bivalent IgG binding buffering of mutations depends on Fab AFFINITY. **(a)** Biophysical model of how bivalent IgG binding buffers mutations that reduce monovalent Fab affinity when the Fab has sufficiently high affinity for the viral antigen, but this buffering vanishes if the Fab affinity is more modest. **(b)** Predictions of a quantitative biophysical model (see Supplementary Appendix) of how a viral mutation that reduces Fab affinity affects both IgG and Fab neutralization of a viral strain to which the Fab has high or low affinity. When the Fab has high affinity, mutations that reduce affinity do not appreciably reduce IgG neutralization. **(c)** Experimental data showing how two known RSV subtype B nirsevimab escape mutations, K68Q and (K/N)201S, affect neutralization by nirsevimab IgG and Fab of pseudoviruses expressing RSV F from subtype A (Long strain) or subtype B (B1 strain). For all plotted neutralization curves, points represent the mean and standard error of at least two replicate measurements. Nirsevimab has a higher affinity for subtype A than subtype B^19,26^. Consistent with the biophysical model, the mutations reduce Fab neutralization for both subtypes, but only reduce IgG neutralization for subtype B.

We then experimentally measured nirsevimab’s neutralization of pseudovirus expressing F from subtype A or B with two previously described nirsevimab escape mutations K68Q and (K/N)201S^20^ (Figure 1c). The F-protein mutations reduced Fab neutralization of both subtypes but only reduced IgG neutralization for subtype B, consistent with the biophysical model (Figure 1b,c). These results suggest that nirsevimab escape mutations are more prevalent in subtype B than A because bivalent IgG binding buffers viral mutations in subtype A due to nirsevimab’s higher affinity for that subtype^19,26^. A corollary, which we build on in this paper, is that it can be more generalizable across viral strains to measure how mutations affect Fab neutralization, since the buffering effect of bivalent binding manifests for IgG but not Fab.

### Functional constraint on F

We used pseudovirus deep mutational scanning^30^ to measure the ability of all amino acid mutants of F to enable pseudotyped lentiviral particles to infect 293T cells expressing the TIM1 attachment factor^31^ (see Methods and Extended Data Figures 1 and 2). These pseudoviruses encode no viral proteins other than F and so can only undergo a single round of cell entry, meaning that they are not fully infectious biological agents capable of causing disease, and so provide a safe way to study mutations to F. We performed the deep mutational scanning using F from the lab-adapted subtype A RSV strain termed “Long” after the name of the patient from whom it was isolated in the 1950s^32^. We created two independent F-expressing pseudovirus libraries, each of which contained nearly all possible amino acid mutations to F’s ectodomain (Extended Data Figure 1). Most F mutants contained just a single amino acid mutation, although some contained no or multiple mutations (Extended Data Figure 1). We quantified the effects of mutations as the log2 cell entry of each mutant relative to the unmutated F, using global epistasis models^33,34^ to jointly analyze the single and multi-mutant data. Negative values indicate a mutation impairs cell entry, while positive values indicate it improves cell entry.

Measurements of how all mutations affect F’s cell entry function are shown in Figure 2a and the interactive plots at https://dms-vep.org/RSV_Long_F_DMS/cell_entry.html. Variants with only synonymous mutations had wildtype-like cell entry scores of zero, variants with stop codon mutations had highly negative scores, and variants with amino acid mutations had scores ranging from wildtype-like to highly negative (Extended Data Figure 2b). We performed three experimental replicates with each of the two independent libraries, and the measured cell entry effects were highly correlated across replicates and libraries (Extended Data Figure 2c,d); we report the average across all replicates for both libraries. We validated that the deep mutational scanning measurements correlated with the infectious titer of pseudoviruses expressing individual F mutations spanning a range of effects (r = 0.92; Figure 2b).

**Figure 2.**
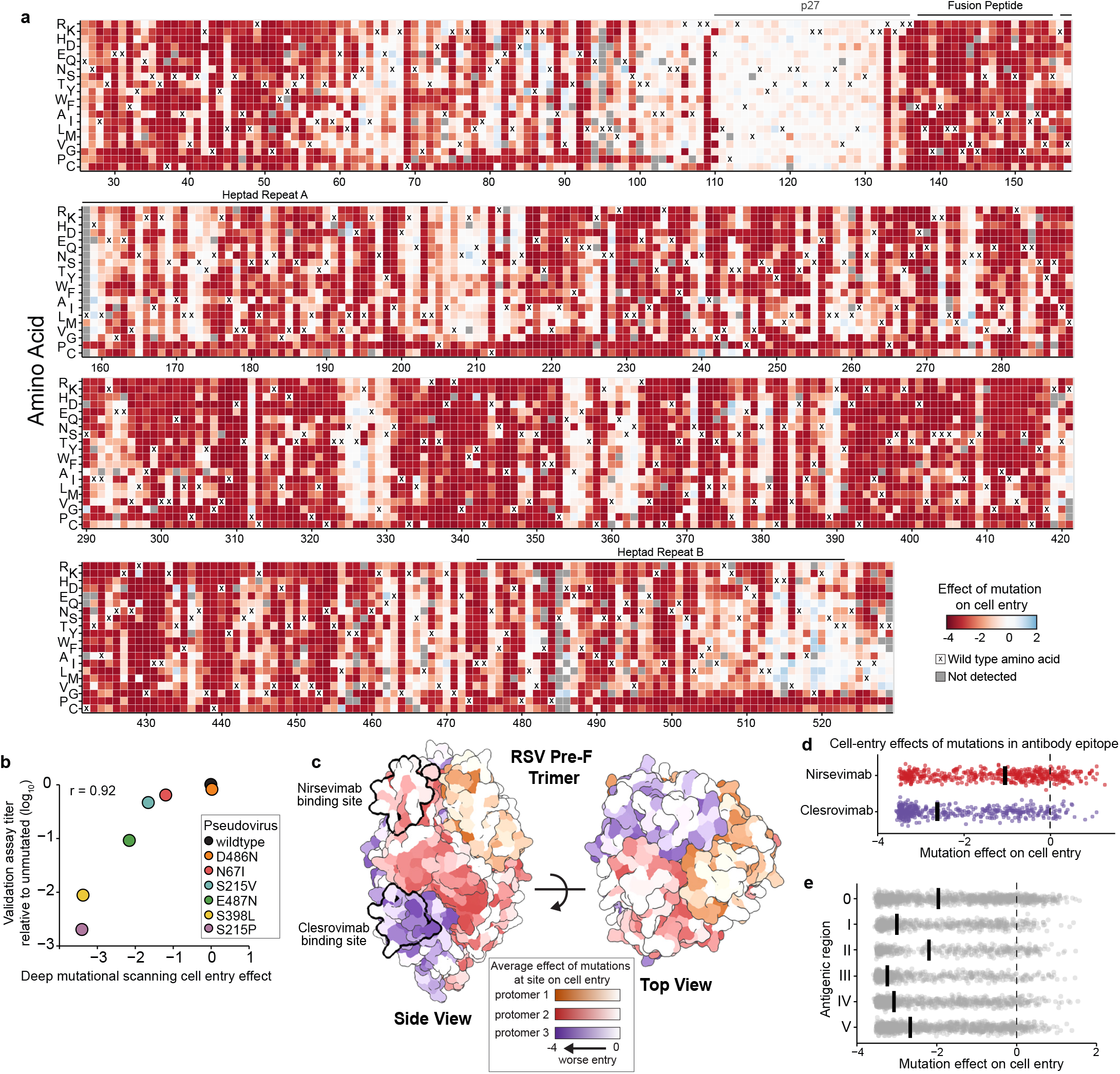
Effects of mutations to RSV F on its cell entry function. **(a)** Effects of single amino acid mutations on pseudovirus entry in 293T-TIM1 cells as measured by deep mutational scanning. Each box represents the effect of a specific mutation, with colors indicating decreased (red), unchanged (white), or slightly increased (blue) cell entry relative to the unmutated F. Mutations not reliably measured in our experiments are indicated by gray boxes, and the wildtype amino acid in the parental subtype A Long strain at each site is indicated with an ‘x’. See https://dms-vep.org/RSV_Long_F_DMS/cell_entry.html for an interactive version of this heatmap. **(b)** Correlation between mutation effects on cell entry measured by deep mutational scanning and pseudovirus titer in a validation assay using F single-mutant pseudoviruses. **(c)** Average effect of mutations at each site mapped onto the prefusion F trimer structure (PDB 5UDC^13^), viewed from the top and side. Each protomer is colored with a separate color with darker shading indicating more deleterious effects on cell entry. The nirsevimab and clesrovimab binding footprints are outlined on one protomer. **(d)** Distribution of mutation effects on cell entry at sites in the epitopes of nirsevimab and clesrovimab (sites that bury ≥5 Å^2^ of surface area upon Fab binding). Each point represents an individual mutation (negative values indicate worse cell entry), and the thick black lines indicate the median. **(e)** Distribution of mutation effects on cell entry for mutations at sites that comprise broadly defined antigenic regions^36^. Thick black lines indicate the median. In panels (d) and (e) the numbers of mutations in each category are: nirsevimab = 473, clesrovimab = 431, site 0 = 1274, site I = 752, site II = 451, site III = 780, site IV = 937; site V = 1324; the points are the mean effects across the three experimental replicates with each duplicate library.

Many regions of F including the fusion peptide are highly constrained with most mutations deleterious for cell entry (Figure 2a). The p27 fragment is cleaved from the mature protein to yield the active form of F; the basic residues comprising the cleavage motifs at each end of p27 are intolerant to mutations whereas the cleaved p27 fragment itself is tolerant to mutations (Figure 2a). The epitope targeted by nirsevimab at the apex of the trimer is more mutationally tolerant than the epitope targeted by clesrovimab on the lateral face of the trimer (Figure 2c,d). Of the broadly defined antigenic regions^35,36^, region 0 is the most mutationally tolerant whereas regions III and IV are more constrained (Figure 2e).

Most F mutations observed in natural RSV sequences relative to the lab-adapted parental strain used in our deep mutational scanning have minimal impact on cell entry (Extended Data Figure 3), as expected since there is strong natural selection for F to retain this function. The main exception is that cell entry is impaired by two mutations (S99N and T101P) that revert sites near an F proteolytic cleavage motif from their identities in the lab-adapted strain to amino acids commonly observed in natural sequences (Extended Data Figure 3). This exception highlights the fact that there can be discrepancies between how mutations affect pseudovirus entry in cell culture versus real-world viral fitness: presumably the N99 and P101 identities in natural sequences are better for authentic viral infection of human airway cells even if S99 and T101 give better pseudovirus entry into a cell line.

### Nirsevimab neutralization of F mutants

We next measured how all functionally tolerated F mutations affect neutralization by nirsevimab, which was the first antibody widely approved for RSV prophylaxis for infants^37^. We incubated the pseudovirus libraries with a range of antibody concentrations and quantified the ability of each variant to infect cells, converting sequencing counts to fraction pseudovirus infectivity at each concentration using a spike-in standard (Extended Data Figure 4a)^30^. We measured the effects of mutations on neutralization by both IgG and Fab forms of nirsevimab, since the biophysical model described above suggests that Fab measurements are more generalizable across genetic backgrounds due to the absence of affinity-dependent buffering of bivalent IgG neutralization. We made two replicate measurements for each of the two independent pseudovirus libraries; throughout we report the average across replicates.

The mutations that affect neutralization by nirsevimab are at a subset of sites from 64 to 73 and 201 to 216 (Figure 3a,b and interactive plots at https://dms-vep.org/RSV_Long_F_DMS/nirsevimab_neutralization.html). These sites are all in or near the structurally determined nirsevimab binding site (Figure 3c). Some mutations at these sites (eg, 64T, 65E, 68Q/N/E, 201S/T, 203I, 208S/D/Y) have been previously shown to affect nirsevimab neutralization^3,5,16–20,22^ or fusion inhibition (eg, 64M/65R, 64V/65E, 68I, 204S, 205S, 208I/K/Y)^4,6^. However, our deep mutational scanning identified numerous mutations that reduce neutralization that have not previously been reported, including mutations to additional amino acid identities at sites 64, 65, 68, 201, 204, 205, 208 and 209 and mutations at sites not previously reported to affect nirsevimab neutralization, including 67, 73, 206-207, 210-211, and 215-216 (Figure 3b).

**Figure 3.**
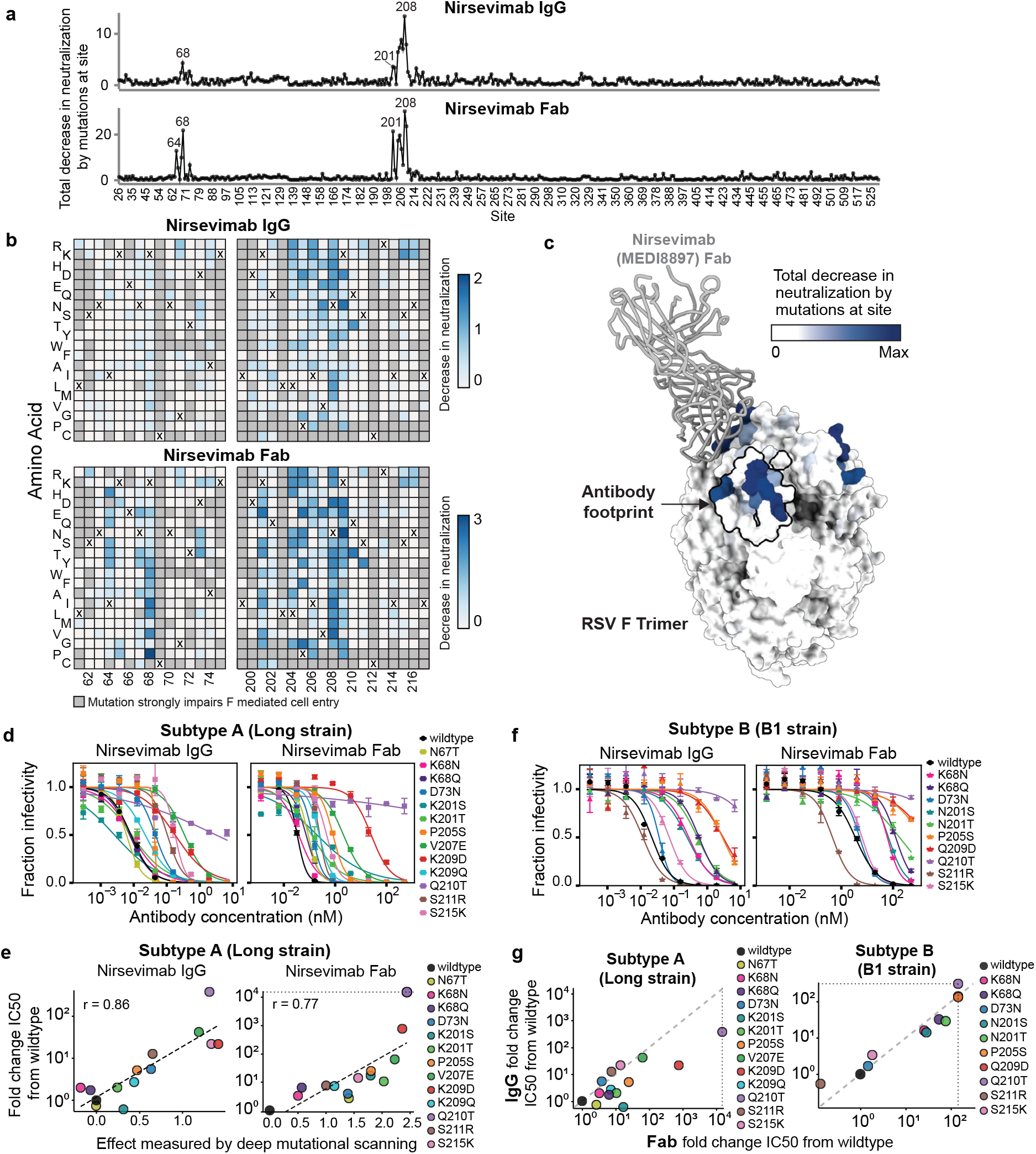
Effects of mutations to RSV F on neutralization by nirsevimab IgG and Fab. **(a)** Total decrease in neutralization from all mutations at each site for nirsevimab IgG or Fab. See https://dms-vep.org/RSV_Long_F_DMS/nirsevimab_neutralization.html for interactive plots that show all mutations. **(b)** Heatmaps showing effects of mutations within the epitope of nirsevimab. Each box represents a single mutation, with positive values indicating decreased neutralization. Dark gray indicates mutations that are too deleterious for cell entry to measure their effect on neutralization. The wildtype amino acid in the subtype A Long strain used for the deep mutational scanning is indicated with an ‘x’. **(c)** Pre-fusion RSV F trimer (PDB 5UDC^13^) in complex with nirsevimab Fab (gray cartoon) colored by the total effect of mutations at each site on neutralization by nirsevimab Fab. **(d)** Neutralization curves for nirsevimab IgG or Fab versus pseudovirus expressing the indicated Long strain F mutants. **(e)** Correlation of mutation effects on neutralization by nirsevimab in the neutralization assays versus deep mutational scanning for IgG or Fab for mutants of the subtype A Long strain F. Horizontal dashed line indicates limit of detection. **(f)** Neutralization curves for nirsevimab IgG or Fab versus pseudovirus expressing the indicated mutants of F from the subtype B strain B1. **(g)** Comparison of fold change in IC50 between nirsevimab IgG and Fab for mutations in subtype A background (Long strain) and subtype B background (B1 strain). In panels (d) and (f), curves for K68Q and K/N201T are duplicated from Figure 1.

To validate the deep mutational scanning, we generated pseudoviruses carrying key mutations in F from the subtype A Long strain and measured their impact in neutralization assays with nirsevimab IgG and Fab (Figure 3d). The effects of mutations on neutralization IC50 were highly correlated with the deep mutational scanning (Figure 3e). Note that these assays validate some of the resistance mutations newly identified by our deep mutational scanning (eg, 73N, 207E, 209D/Q, 210T, 211R, 215K).

To test if our deep mutational scanning of how mutations affect nirsevimab Fab neutralization of F from a subtype A strain could be extrapolated to a subtype B strain, we generated pseudoviruses carrying key mutations in the F of a subtype B strain (B1, a lab-adapted strain from the 1980s^38^). Most mutations that reduced Fab neutralization of the subtype A strain also reduced both Fab and IgG neutralization of the subtype B strain (Figure 3f and Extended Data Figure 4b,c). Consistent with the biophysical model, deep mutational scanning measurements of how subtype A mutations affect Fab neutralization correlate more strongly with both IgG and Fab neutralization of subtype B mutants than do deep mutational scanning measurements performed with IgG (Extended Data Figure 4c). The biophysical model also predicts that F mutations will cause a greater reduction in Fab than IgG neutralization for subtype A, but affect IgG and Fab neutralization similarly for subtype B since there is bivalent IgG buffering for the higher-affinity subtype A binding but not the lower-affinity subtype B binding; our measurements validate this prediction (Figure 3g).

Note that S211R slightly increases subtype B neutralization despite decreasing subtype A neutralization for both Fab and IgG; for this mutation there must be additional background-specific differences between subtypes.

Our deep mutational scanning also identifies the F sequence differences responsible for the lower nirsevimab Fab potency to subtype B than subtype A. Most subtype A strains have N at site 67 and K at 209, while most subtype B strains have T at 67 and Q or R at 209. In the deep mutational scanning, N67T and K209Q reduce neutralization by nirsevimab Fab. We confirmed that mutating the subtype A F to the subtype B identities at these sites (N67T and K209Q) reduces neutralization by nirsevimab Fab, while the reverse mutations in subtype B (T67N and Q209K) increase neutralization (Extended Data Figure 4d). The effects of these swap mutations are much larger on Fab versus IgG neutralization, consistent with the bio-physical model which posits that the nirsevimab affinity for F is sufficiently high that bivalent binding largely buffers mutation effects on IgG neutralization (Extended Data Figure 4d).

Measuring mutation effects on nirsevimab Fab neutralization identifies F mutations to subtype A that impact IgG neutralization in combination even when they individually have little effect. Specifically, the F mutations N67T, K68Q, K201S, and K209Q individually reduce nirsevimab Fab but not IgG neutralization of subtype A— but reduce IgG neutralization when combined as double or triple mutants (Extended Data Figure 5). These results show the relevance of characterizing mutational impacts on Fab neutralization for understanding IgG neutralization of multiply mutated F variants.

### Clesrovimab neutralization of F mutants

Clesrovimab, which targets a different region of F than nirsevimab, was recently approved as another option for RSV prevention for infants^39,40^. We measured the effects of F mutations on neutralization by clesrovimab IgG and Fab, performing two replicate deep mutational scanning experiments for each of the two independent pseudovirus libraries.

The mutations that most affect clesrovimab neutralization occur at a subset of sites between 426-470 (Figure 4a,b). These sites are within or near the structurally determined clesrovimab binding site (Figure 4c and interactive plots at https://dms-vep.org/RSV_Long_F_DMS/clesrovimab_neutralization.html). Some mutations identified by our deep mutational scanning have been previously reported to affect clesrovimab (e.g., 443P, 445N, and 446E/R/W^14,40^). But we also identified new mutations affecting clesrovimab, including additional mutations at sites 443, 445, and 446, as well as mutations at sites not previously associated with clesrovimab resistance (426, 429, 433, and 470) (Figure 4b).

**Figure 4.**
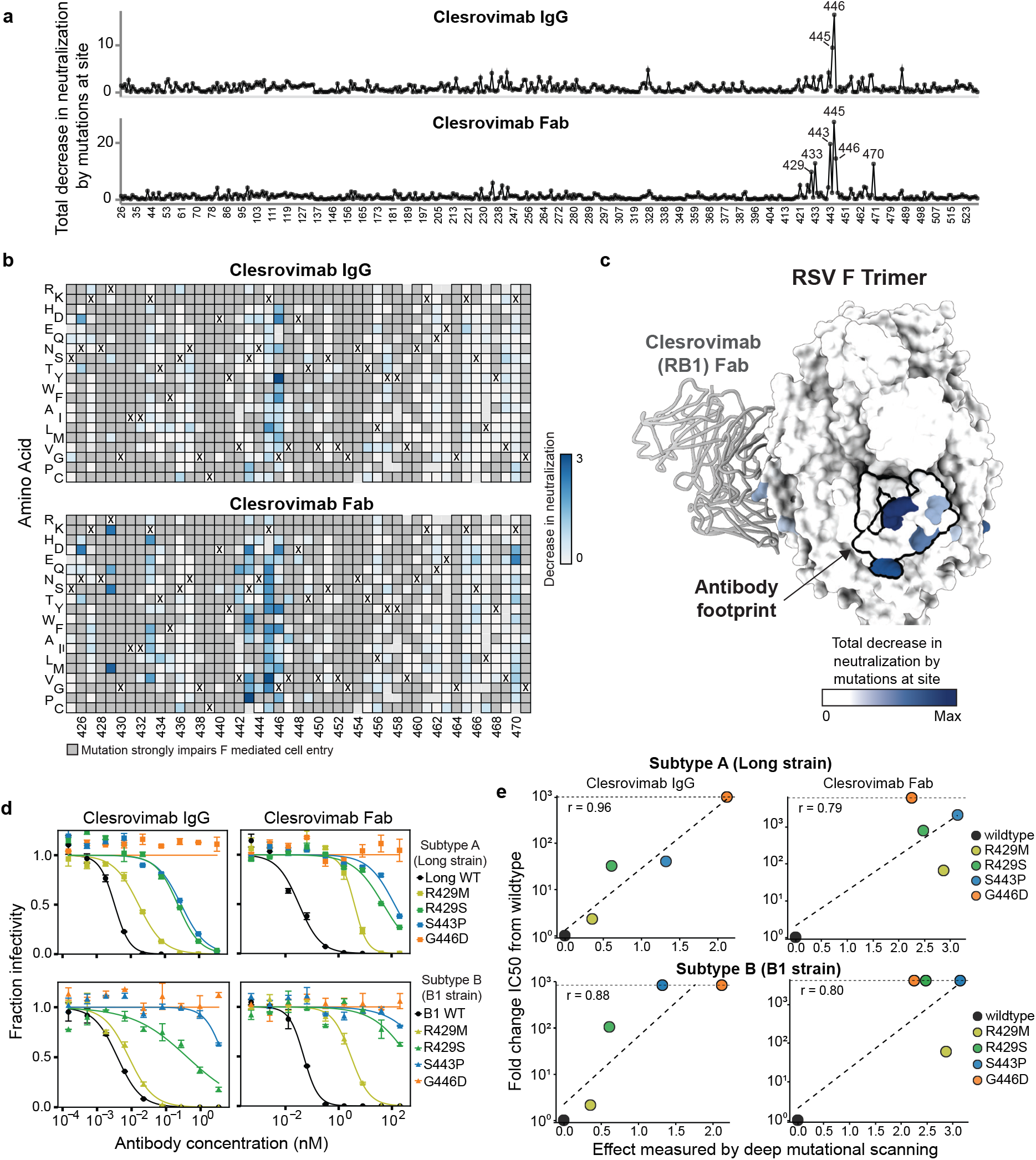
Effects of mutations to RSV F on neutralization by clesrovimab IgG and Fab. **(a)** Total decrease in neutralization from all mutations at each site for clesrovimab IgG or Fab as measured by deep mutational scanning. See https://dms-vep.org/RSV_Long_F_DMS/clesrovimab_neutralization.html for interactive plots that show the effects of all mutations. **(b)** Heatmaps showing effects of mutations within the epitope of clesrovimab. Each box represents a single mutation, with positive values indicating decreased neutralization. Dark gray indicates mutations that are too deleterious for cell entry to measure their effect on neutralization. Light gray shading indicates mutations that were not measured. The wildtype amino acid in the subtype A Long strain used for the deep mutational scanning at each site is indicated with an ‘x’. **(c)** Pre-fusion RSV F trimer (PDB 6OUS^14^) in complex with clesrovimab Fab (gray cartoon) colored by the total effect of mutations at each site on neutralization by clesrovimab Fab. **(d)** Neutralization curves for clesrovimab IgG and Fab of pseudovirus expressing F with point mutations in subtype A (Long) or subtype B (B1) background. Individual mutations were selected to span a range of effects on neutralization. **(e)** Correlation of mutation effects on neutralization by clesrovimab IgG and Fab between deep mutational scanning and traditional pseudovirus neutralization assays with the indicated F mutants in subtype A (Long strain) or subtype B (B1 strain). Horizontal and vertical lines indicate limits of detection.

To validate the deep mutational scanning, we generated pseudoviruses carrying key mutations in F from both subtype A (Long strain) and subtype B (B1 strain) and measured their neutralization by clesrovimab IgG and Fab (Figure 4d,e). The effects of mutations on neutralization IC50 were highly correlated with the deep mutational scanning for both subtypes (Figure 4e).

Mutations that moderately reduce clesrovimab neutralization had a greater impact on Fab than IgG neutralization for both subtypes (Figure 4a,b,d; for example, R429M/S). This tendency for moderate-effect mutations to reduce Fab neutralization more than IgG neutralization is also observed for nirsevimab against subtype A but not subtype B (Figure 3g). These observations are consistent with the biophysical model of how bivalent IgG buffering reduces the impact of mutations on IgG neutralization when the Fab affinity is sufficiently high (which is true for clesrovimab against both sub-types A and B, but for nirsevimab only against subtype A). Unlike for nirsevimab, clesrovimab Fab has similarly high neutralization potencies against both subtypes (Extended Data Figure 6a), and the effects of mutations on clesrovimab neutralization are well correlated across subtypes (Extended Data Figure 6b).

Our deep mutational scanning found fewer sites where mutations strongly reduced clesrovimab neutralization compared to nirsevimab. Part of the reason may be differences in functional constraint: it is only possible to measure the impact on neutralization of F mutations that retain at least some cell entry function, and mutations in the clesrovimab epitope tend to be more deleterious to cell entry than mutations in the nirsevimab epitope (Figure 2c,d). Indeed, even among mutations that retain sufficient cell entry to measure their impact on neutralization, the ones that reduce clesrovimab neutralization tend to impair cell entry more than the ones that reduce nirsevimab neutralization (Extended Data Figure 6c). However, there are still some mutations that reduce clesrovimab neutralization without impairing cell entry.

### Phenotypically-informed RSV surveillance

Now that nirsevimab and clesrovimab are in widespread use, surveillance for natural RSV strains with resistance to these antibodies is a public health priority^3,4,6^. Although large numbers of human RSV infections are regularly being sequenced^3–6,20,41–44^, interpretation of these sequences has been limited by incomplete knowledge of which mutations affect antibody resistance. Our deep mutational scanning enables immediate assessment of all RSV sequences for antibody resistance. We calculated two escape scores for each RSV sequence: the summed effect on neutralization of all its F mutations relative to the subtype A Long strain used in the deep mutational scanning, and the effect of the single mutation that caused the greatest reduction in neutralization. We integrated these scores into Nextstrain^45^ phylogenetic trees of RSV showing the latest sequence data from Pathoplexus^46^. Since Nextstrain subsamples the many available F sequences for effective visualization, we also created new builds that ensure the subsampling includes sequences with high escape scores. Interactive phylogenetic trees that can be colored by the escape scores and are updated to include the latest available sequences are at https://nextstrain.org/rsv/b/F-antibody-escape/6y?c=Nirsevimab-Fab_total_escape. Static images based on further subsampled versions of these trees are in Figure 5a,b and Extended Data Figure 7a,b.

**Figure 5.**
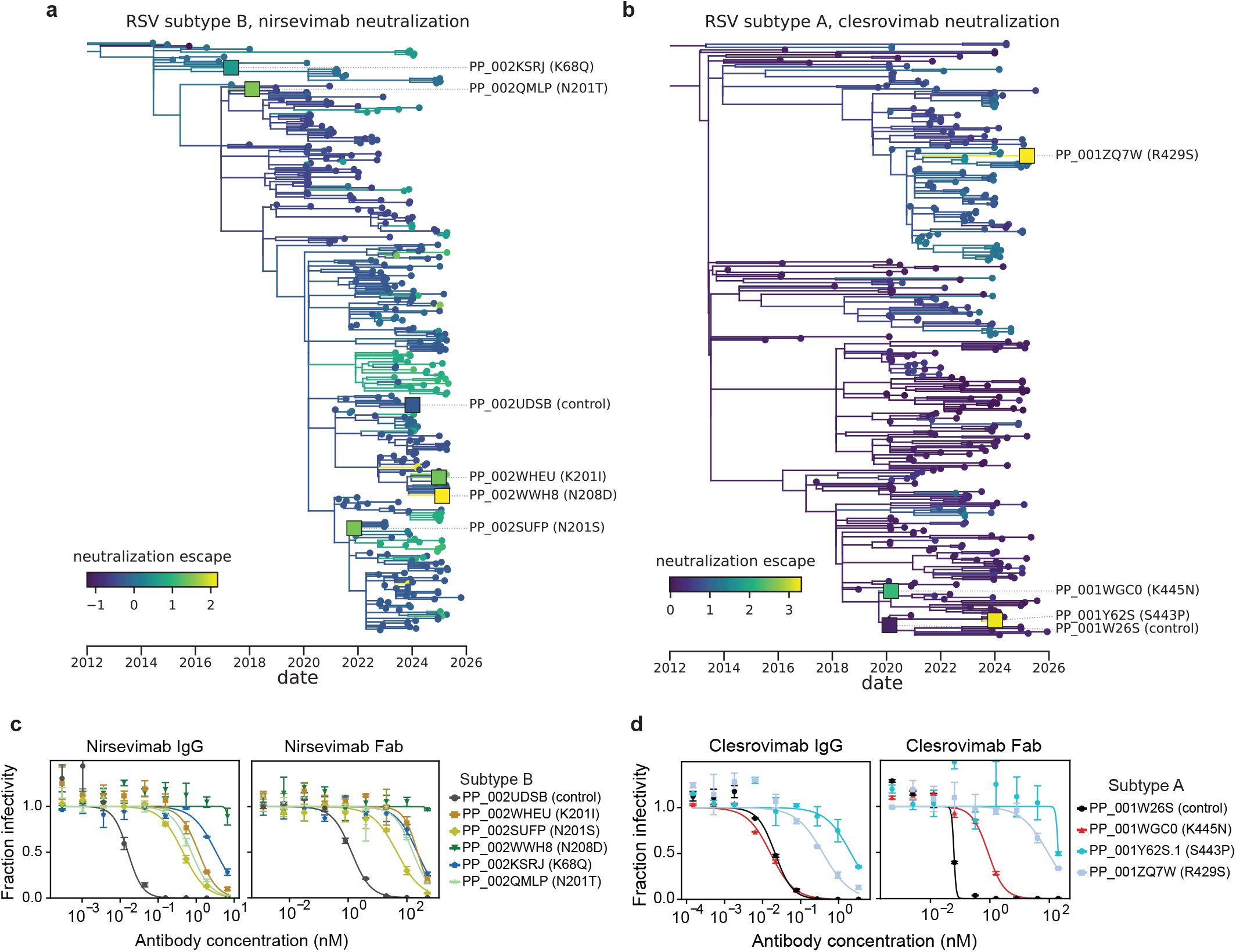
Sporadic natural RSV strains have reduced neutralization by nirsevimab or clesrovimab. **(a)** Phylogenetic trees of subtype B RSV F sequences colored by nirsevimab escape scores computed as the summed effects of all mutations as measured by the deep mutational scanning. Strains chosen for validation of nirsevimab neutralization are indicated by boxes and labeled with the Pathoplexus identifier and top resistance mutation. See https://nextstrain.org/rsv/b/F-antibody-escape/6y?c=Nirsevimab-Fab_total_escape for interactive Nextstrain trees that show more sequences and are updated in real time. **(b)** Tree of subtype A RSV F sequences colored by clesrovimab escape scores https://nextstrain.org/rsv/a/F-antibody-escape/6y?c=Clesrovimab-Fab_total_escape. **(c)** Neutralization curves showing nirsevimab IgG or Fab neutralization of pseudoviruses expressing F from the subtype B strains labeled on the tree. The strains with high escape scores all have reduced neutralization relative to a control strain with a low escape score. **(d)** Neutralization curves showing clesrovimab IgG or Fab neutralization of pseudoviruses expressing F from the subtype A strains labeled on the tree. Strains with high escape scores all have reduced neutralization relative to a control strain, except a strain (PP_001WGC0) with only a moderate escape score that shows reduced Fab but not IgG neutralization. See Extended Data Figure 7 for trees and validating neutralization assays for subtype A strains with nirsevimab resistance and subtype B strains with clesrovimab resistance.

Sequences with high nirsevimab or clesrovimab escape scores were rare (<1% of all sequences) and distributed across the phylogenetic trees, suggesting resistance mutations have arisen sporadically and not undergone sustained spread (Figure 5a,b, Extended Data Figure 7a,b, and https://nextstrain.org/rsv/b/F-antibody-escape/6y?c=Nirsevimab-Fab_total_escape). However, the fact that strains with resistance have emerged and transmitted in humans to a limited extent underscores the importance of continued surveillance.

To validate that natural sequences with high escape scores had reduced neutralization, we generated RSV pseudoviruses expressing the F proteins from strains with high escape scores for nirsevimab or clesrovimab. Nearly all strains with high escape scores had reduced neutralization relative to control recent strains with low escape scores (Figure 5c,d and Extended Data Figure 7c,d). The natural strains with reduced neutralization include ones with resistance mutations newly identified by our deep mutational scanning (eg, K201I and K201E for nirsevimab, and R429S for clesrovimab). A caveat is that the escape scores assume resistance can be predicted from the additive effects of a strain’s mutations; our validation assays show that assumption usually holds, but we did identify one subtype A strain with a high nirsevimab escape score that did not have reduced neutralization, possibly due to an epistatic interaction of its S211R resistance mutation with a nearby R213S mutation (Extended Data Figure 7c).

### Historical and candidate antibodies

We next used deep mutational scanning to characterize mutations that affect neutralization by five additional antibodies of historical or potential future clinical relevance.

Suptavumab failed a Phase 3 trial run from 2015 to 2017 due to lack of efficacy against subtype B^2^. Our deep mutational scanning shows that the mutations that most reduce suptavumab neutralization are at sites 173 and 174 (Figure 6a-c). Starting in 2015, L172Q and S173L became prevalent in subtype B sequences (Figure 6d), and the reduced ability of suptavumab to neutralize strains with these two mutations led to the failed clinical trial^2^. Our deep mutational scanning shows that S173L reduces suptavumab neutralization while having minimal adverse impact on F’s cell entry function (Figure 6c). Notably, S173L had already spread to roughly half of subtype B sequences by the time the suptavumab Phase 3 clinical trial began in November of 2015^2^, so suptavumab’s failure could have been anticipated if our deep mutational scanning had been available in advance.

**Figure 6.**
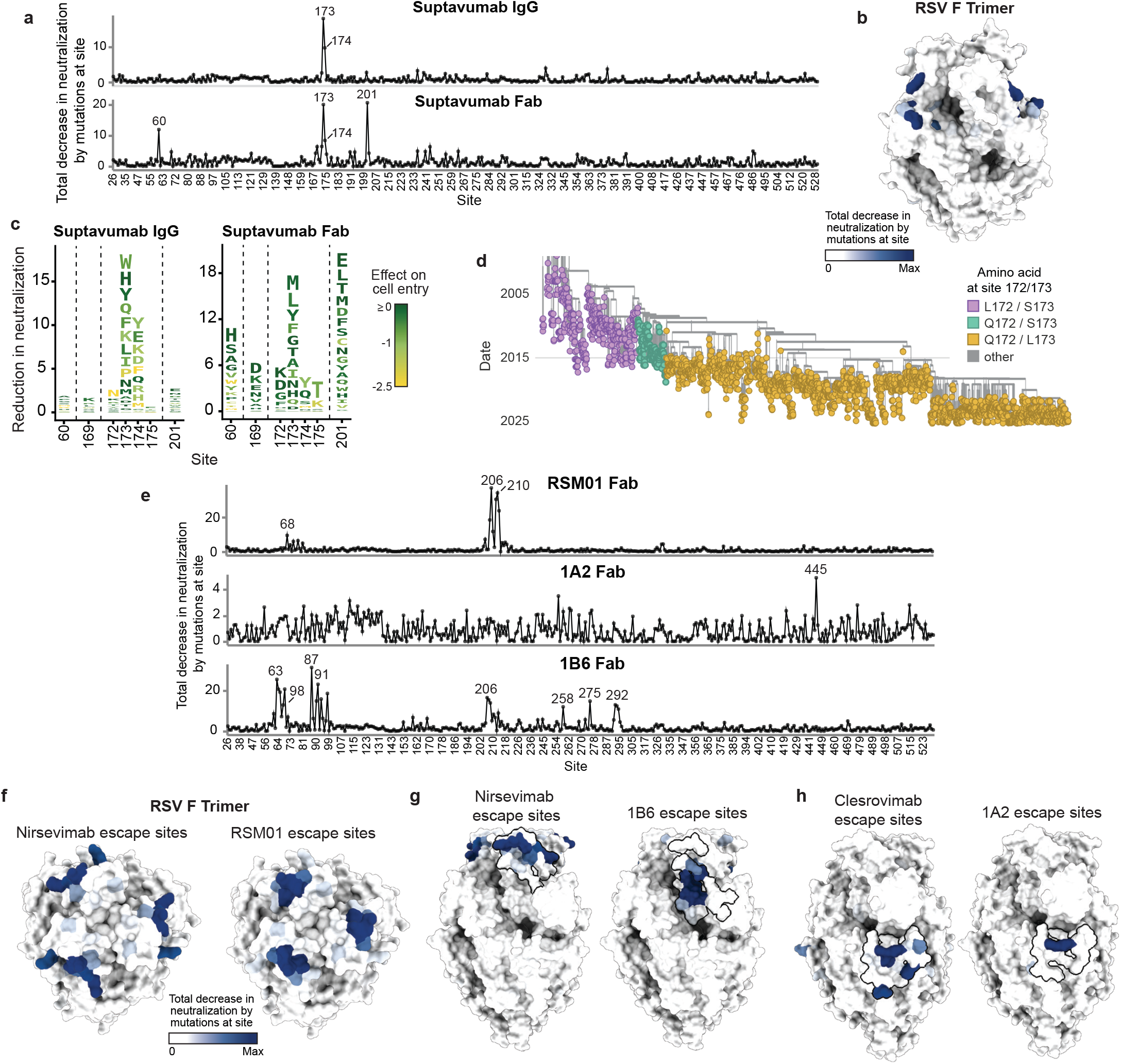
Effects of mutations to RSV F on neutralization by the historical failed antibody suptavumab and three candidate future clinical antibodies (RSM01, 1A2, 1B6) **(a)** Total decrease in neutralization from all mutations at each site for suptavumab IgG or Fab as measured by deep mutational scanning. See https://dms-vep.org/RSV_Long_F_DMS/suptavumab_neutralization.html for interactive plots that show the effects of all mutations. **(b)** Pre-fusion RSV F trimer (PDB 5UDC^13^) colored by the total effect of mutations at each site on neutralization by suptavumab Fab. **(c)** Effects on suptavumab neutralization of mutations at key sites. The height of each letter is proportional to the reduction in neutralization caused by a mutation to that amino acid. Mutations are colored by their effect on F-mediated cell entry in the absence of antibody (dark green indicates no effect on cell entry and yellow indicates reduced cell entry). **(d)** Phylogenetic trees of subtype B RSV F sequences colored by amino acid identities at sites 172 and 173. **(e)** Total decrease in neutralization from all mutations at each site for RSM01, 1A2, or 1B6 Fabs. See https://dms-vep.org/RSV_Long_F_DMS/ for interactive plots that show effects of all mutations. **(f)** Pre-fusion RSV F trimer (PDB 5UDC^13^) colored by the total effect of mutations at each site on neutralization by RSM01 or nirsevimab Fab. **(g)** Pre-fusion RSV F trimer (PDB 9LLY^52^) colored by the total effect of mutations at each site on neutralization by 1B6 or nirsevimab Fab. The structurally determined antibody-binding footprint is outlined. **(h)** Pre-fusion RSV F trimer (PDB 9LLY^52^) colored by the total effect of mutations at each site on neutralization by clesrovimab or 1A2.

Palivizumab is an older monoclonal antibody previously recommended only for high-risk (eg, pre-term) infants^47^ that has now largely been replaced by nirsevimab and clesrovimab since they have a longer half-life and are more potent. Our deep mutational scanning showed palivizumab neutralization is strongly reduced by mutations at a subset of sites from 255 to 277 (Extended Data Figure 8), consistent with prior work showing mutations at some of these sites cause palivizumab resistance48–51.

We next characterized three anti-RSV antibodies that are candidates for_1 0_future clinical use. RSM01 has been suggested for use in low- and middle-income countries where RSV is a leading cause of infant mortality^11,22^, whereas 1A2 and 1B6 target different epitopes on F and have been proposed as components of an antibody cocktail^52^. The IgGs of all three antibodies potently neutralized pseudoviruses with either a subtype A or B F with IC50s less than 0.1 nM, as do the existing clinical antibodies clesrovimab and nirsevimab (Extended Data Figure 9a). However, the Fabs of the apex-targeting antibodies (nirsevimab, RSM01 and 1B6) have substantially differing potencies between subtypes: nirsevimab and RSM01 Fabs were more potent against subtype A than B, whereas 1B6 Fab was more potent against subtype B than A (Extended Data Figure 9a). The Fabs of the lateral-face targeting antibodies clesrovimab and 1A2 were similarly potent against both subtypes (Extended Data Figure 9a). Notably, Fab and IgG of 1A2 had similarly high neutralization potencies (Extended Data Figure 9a); likely because this antibody’s exceptionally high affinity for F^52^ means bivalent IgG buffering confers no further neutralization advantage over the Fab, although under the bio-physical model presented earlier in this paper, it is still expected to confer increased IgG robustness to F mutations.

Deep mutational scanning showed that mutations reducing RSM01 neutralization occur at a subset of sites spanning residues 68-80 and 202-216 (Figure 6e,f). Although RSM01 targets a similar region of prefusion F as nirsevimab, there is only partial overlap of the mutations that impact neutralization by these antibodies. For example, mutations at sites 201, 204 and 208 have substantially larger effects on nirsevimab than RSM01 neutralization (Figure 6e,f, Figure 3a,b, and Extended Data Figure 9b). Consistent with these results, a recent study that serially passaged replication-competent RSV in the presence of RSM01 selected an I206T/N262Y double mutant with partial antibody resistance^22^, and our deep mutational scanning identifies I206T as reducing RSM01 neutralization.

The two antibodies comprising the candidate antibody cocktail were impacted by largely distinct sets of mutations. The mutations that reduce neutralization by 1B6 are near the apex of F, but at largely distinct sites from where mutations affect the other apex-targeting antibodies nirsevimab and RSM01 (Figure 6e,g and Extended Data Figure 9b). The sites where mutations affect 1A2 neutralization substantially overlap with sites that affect clesrovimab neutralization, although there are few single mutations that strongly reduce 1A2 neutralization likely because its exceptionally high affinity for F confers substantial robustness to mutations (Figure 6e,h and Extended Data Figure 9c). The mutations that reduce neutralization by all three apex-targeting antibodies (nirsevimab, RSM01, and 1B6) are often better tolerated for F’s cell entry function than mutations that reduce neutralization by the lateral-face targeting antibodies (clesrovimab and 1A2) (Extended Data Figure 9b,c).

## Discussion

We have used pseudovirus deep mutational scanning to measure how nearly all mutations to RSV F affect neutralization by clinically relevant antibodies. Prior work to identify resistance mutations, including for FDA-required resistance analyses^19,40,53^, has serially passaged authentic RSV in the presence of antibody^5,14,22^. Our strategy has two advantages over this classical approach: it characterizes all mutations rather than just identifying whichever ones stochastically arise during viral passage, and it does not generate mutants of actual replicative virus (pseudoviruses can only undergo a single round of cell entry). On the other hand, serial passaging can uncover resistance mediated by combinations of mutations, whereas deep mutational scanning only measures the effects of single mutations. However, in practice our deep mutational scanning identified all resistance mutations previously reported by serial-passage studies as well as a number of new mutations, some of which are present in natural RSV sequences.

At first blush, the fact that we measured effects of mutations to F from only a single subtype A strain might seem a major limitation, especially given the perplexing prior observation that nirsevimab resistance is more prevalent in subtype B^3–6,16–20^. But a key insight of our work is that subtype-dependent effects of mutations on nirsevimab neutralization are largely explained by bio-physically modeling how bivalent IgG binding buffers mutations when Fab potency is sufficiently high. Because nirsevimab Fab has higher affinity for subtype A than subtype B RSV F^19,26^, mutations that moderately reduce Fab neutralization of both subtypes only reduce IgG neutralization of subtype B. Therefore, we measured how F mutations affected both IgG and Fab neutralization, and showed that the Fab measurements identified mutations that reduced IgG neutralization of subtype B despite having minimal impact on IgG neutralization of subtype A. This biophysical principle extends beyond RSV and nirsevimab, and implies that mutations that modulate Fab potency without affecting IgG neutralization can nonetheless alter the impact of subsequent mutations on IgG neutralization. However, other more specific mechanisms can also shape how a mutation impacts different strains, and so the fact that our measurements were performed in a single genetic background remains a caveat.

An immediate benefit of our work is enabling real-time assessment of natural RSV sequences for antibody resistance. Because anti-RSV antibodies are of such public-health importance, genomic surveillance of RSV has become a major priority^4,6,20,21^: there are now >60,000 sequences in public databases^46^ and the number is rapidly growing. However, interpretation of these sequences has been limited by the fact that prior experimental work only characterized how a small fraction of all F mutations affect antibody neutralization^3–6,16–20,22,40^. We used our deep mutational scanning data to score all available sequences and identify sporadic nirsevimab or clesrovimab resistance, in some cases mediated by mutations not previously reported to impact those antibodies. We implemented this resistance-scoring system into real-time Nextstrain phylogenetic trees and created a web interface with our experimental data (https://dms-vep.org/RSV_Long_F_DMS). These resources will enable continued assessment for resistance to current clinical antibodies as well as new ones under development like RSM01, 1B6, and 1A2^11,22,52^.

Fortunately, resistance to nirsevimab and clesrovimab is currently rare. Resistance could remain rare, or could eventually spread as happened with suptavumab resistance in RSV subtype B in 2015-2016^2^ or oseltamivir (Tamiflu) resistance in H1N1 influenza in 2008^54^. Resistance can spread incidentally if resistance mutations also enhance viral fitness by an unrelated mechanism like increasing transmissibility or reducing recognition by population immunity^55,56^; such incidental spread likely explains why suptavumab resistance emerged prior to Phase 3 clinical trials of that antibody. We previously showed that some nirsevimab resistance mutations mildly reduce neutralization of RSV by polyclonal human sera^31^, raising the possibility that they could confer a small incidental benefit to the virus if they do not adversely impact any other aspects of fitness. In addition, now that nirsevimab and clesrovimab are in clinical use, there is also direct evolutionary pressure for viral resistance that enables infection of treated individuals. However the infants receiving these antibodies are just a small fraction (<1%) of all people^57,58^. Resistance would therefore confer a real but small advantage, and so such a strain would spread only if the resistance mutations do not impair other aspects of transmissibility by even a small amount^56^. It is impossible to experimentally measure RSV transmissibility with such precision; although our deep mutational scanning measured how mutations affect F’s cell entry function, these experiments have a precision far worse than 1% and cell entry is only one of several F properties that contribute to transmissibility. Therefore, the best that can currently be done is to monitor for the real-world spread of resistance while continuing to develop new antibodies with distinct resistance profiles—both goals that our study helps advance.

## Extended Data Figures and Legends

**Extended Data Figure 1.**
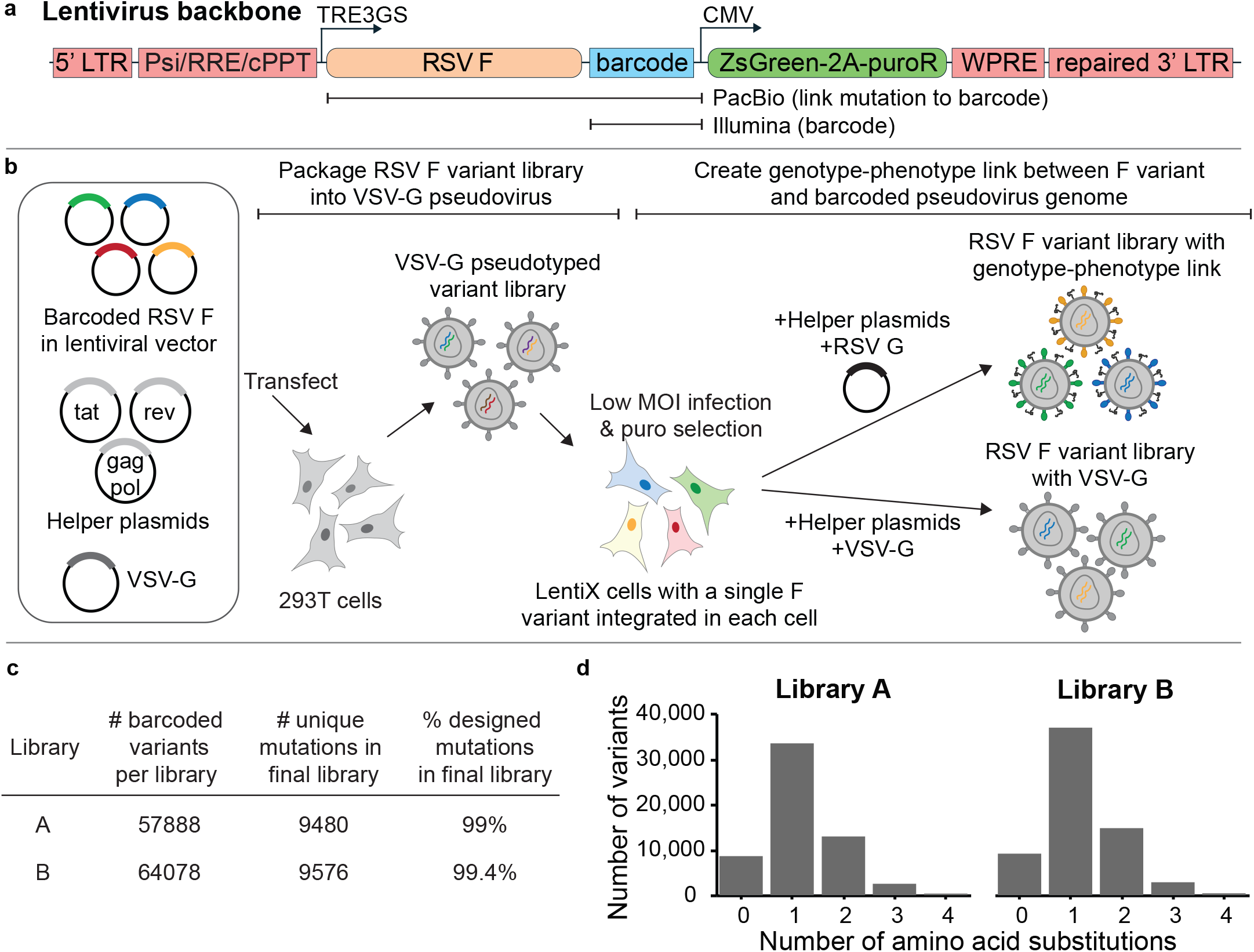
Overview of RSV F deep mutational scanning. **(a)** Lentivirus genome used to make genotype-phenotype linked pseudovirus libraries. Long-read PacBio sequencing is used to link each barcode to a F mutant. Short-read Illumina sequencing is used to quantify the frequency of F variants in experiments. **(b)** Process to generate pseudovirus libraries. We produce VSV-G-pseudotyped lentiviral particles encoding F in their genomes. LentiX cells are transduced by these particles at low MOI (multiplicity of infection, <0.01) so most infected cells receive only one genome integration. Cells with integrated proviruses are selected using puromycin. The library of F-pseudotyped particles is produced by transfecting lentiviral helper plasmids and RSV G. VSV-G pseudovirus is produced as a control to measure library composition independent of F function. PacBio sequencing is performed after integration into cells to account for lentiviral recombination. **(c)** Number of barcoded variants, number of unique F amino acid mutations represented at least once among these variants, and percent of all possible F ectodomain mutations in each of the two pseudovirus libraries. **(d)** Distribution of number of mutations per F variant.

**Extended Data Figure 2.**
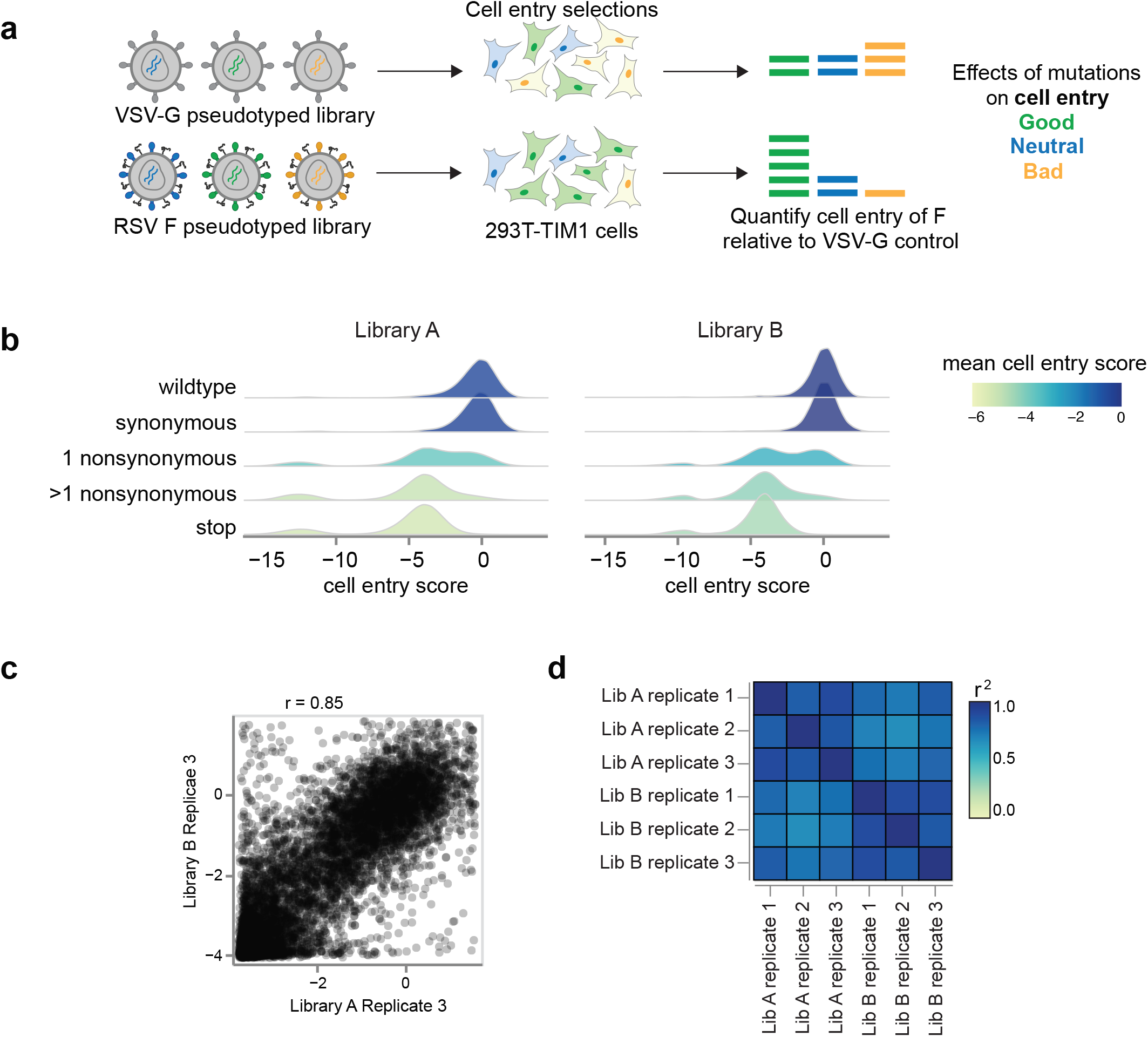
Measurement of how mutations affect F-mediated cell entry. **(a)** Workflow for measuring effects of mutations on cell entry. Pseudovirus libraries expressing RSV F and G or VSV-G are used to infect 293T-TIM1 cells. After infection, lentivirus genomes are isolated from infected cells and barcodes are sequenced to quantify the ability of each F variant to infect cells. Cell entry is quantified as the log2 ratio of the counts of the barcode in the F-mediated entry versus VSV-G control experiments after normalizing these counts to those for the unmutated F protein. So negative values indicate impaired entry and values of zero indicate no effect. **(b)** Distributions of cell entry scores for the two replicate libraries. Histograms are separated by the type of codon mutation in the mutant. **(c)** Pearson correlation (r) of the effect of mutations on cell entry between a single experimental measurement made using each of the two independent libraries. **(d)** Three experimental replicates were done for each of the two libraries. Correlations of the effects of mutations are shown for each pair of replicates.

**Extended Data Figure 3.**
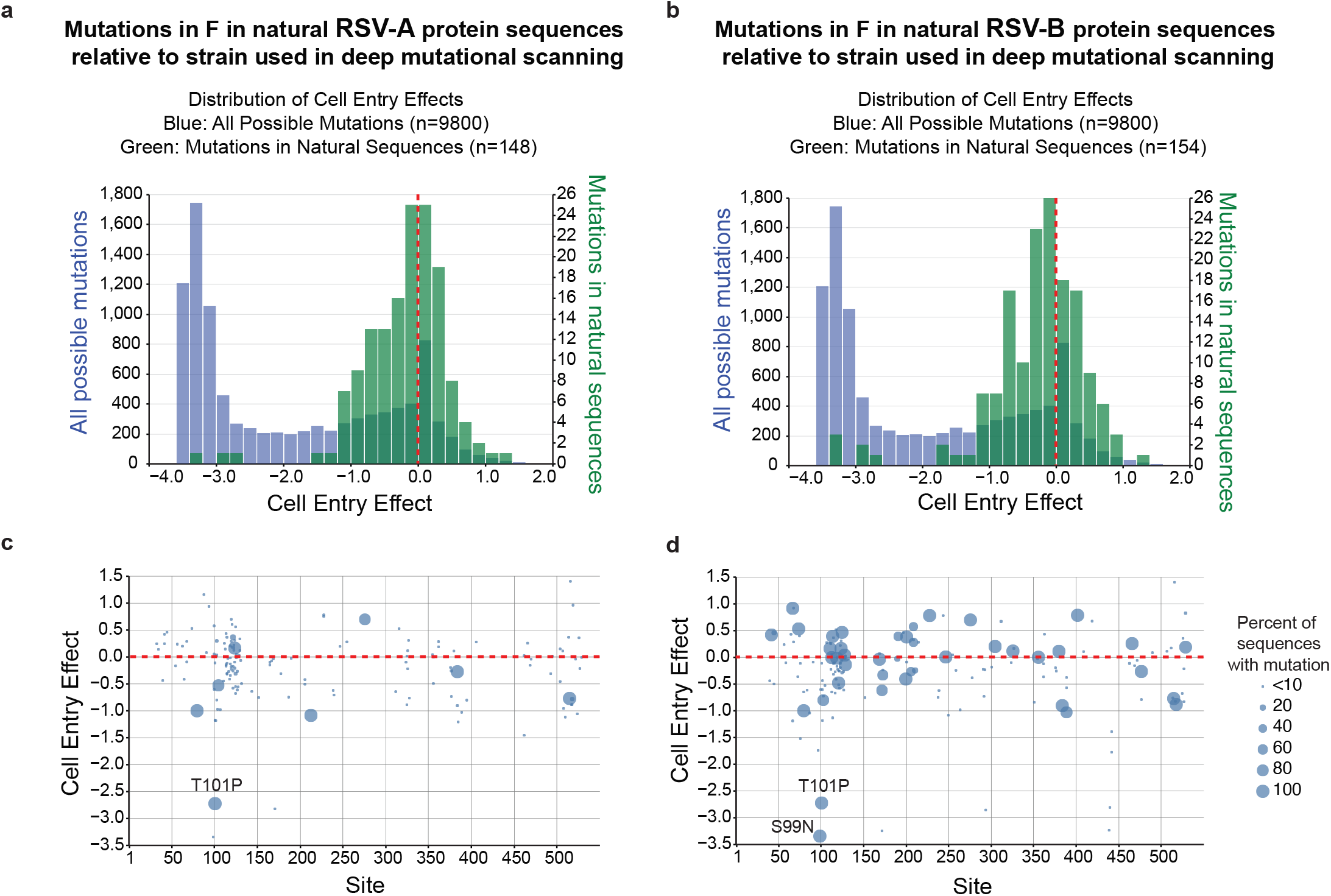
Amino acid differences in natural RSV sequences relative to the strain used for deep mutational scanning mostly have minimal impact on cell entry. **(a**,**b)** Distribution of effects on cell entry (as measured by deep mutational scanning) for all possible F protein mutations (blue) and mutations observed in natural RSV F sequences (green) for subtypes A and B. Mutations are defined relative to the lab-adapted subtype A Long strain used in the deep mutational scanning. The red vertical dashed line indicates an effect of zero, corresponding to no impact on cell entry. **(c**,**d)** Cell entry effects for mutations in natural subtype A and B F sequences relative to the strain used in deep mutational scanning. Each circle represents a mutation, with its x-axis location indicating the site in the protein and its size indicating the percent of F sequences with that mutation. The red dashed line indicates zero effect (neutral). Only two mutations measured to be deleterious to cell entry are at high frequency in natural sequences, T101P and S99N. T101P is found in nearly all natural sequences and S99N is found in nearly all natural subtype B sequences. S99N is likely deleterious in the lab-adapted strain as it creates an N-linked glycosylation site near the F cleavage site. However, this glycosylation site is not formed in natural sequences due to no S/T residues at site 101. T101P is a determinant of human metapneumovirus growth in cell culture^59^, suggesting the difference between the lab-adapted strain and natural sequences may be a lab-adaptation.

**Extended Data Figure 4.**
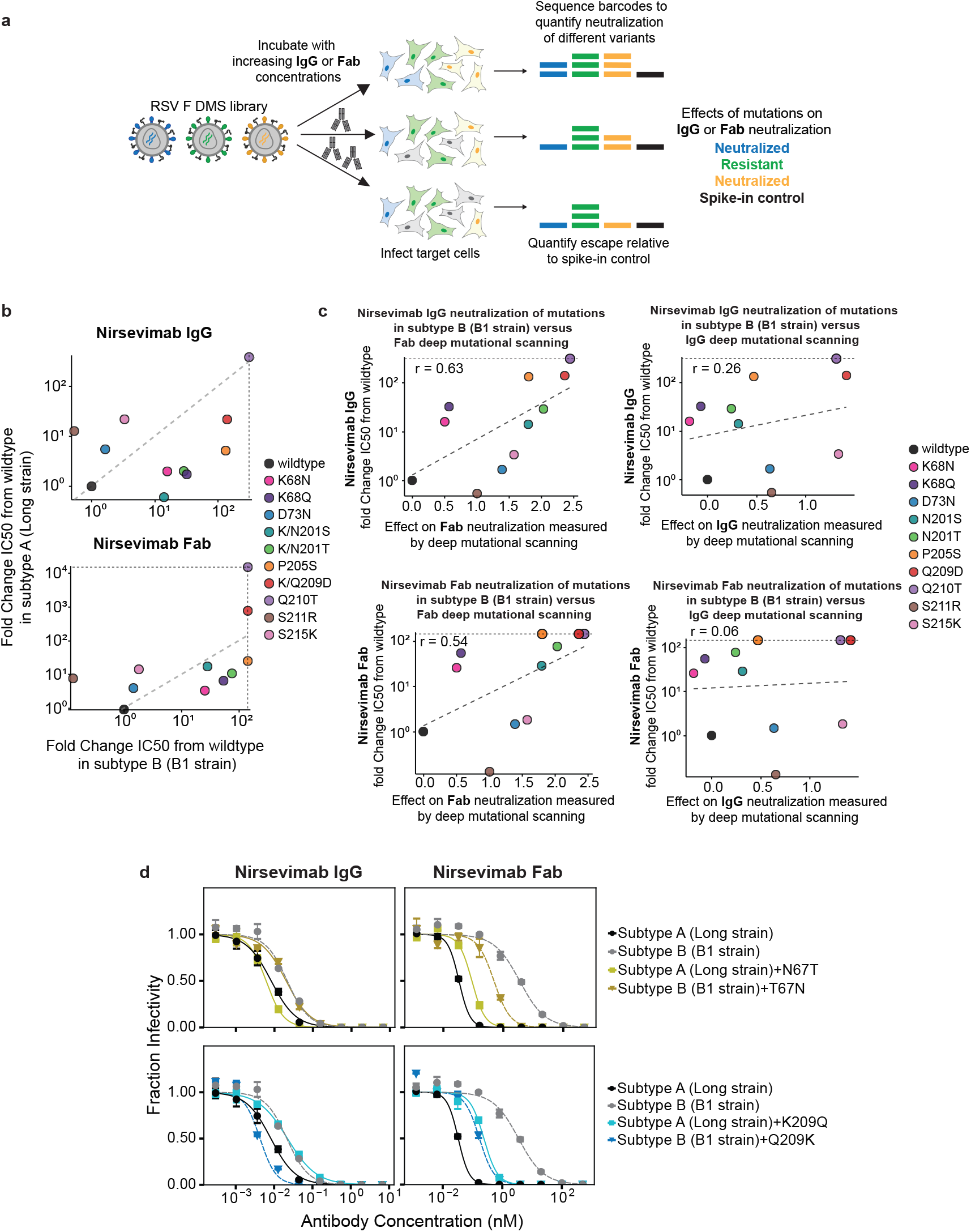
Deep mutational scanning measurements versus effects of mutations on subtype B as measured in validation neutralization assays. **(a)** Workflow for measuring effects of mutations on antibody neutralization using deep mutational scanning. Pseudovirus libraries expressing RSV F and G are used to infect 293T-TIM1 cells after incubation with increasing concentrations of antibody (IgG or Fab). After infection, lentivirus genomes are isolated from infected cells and combined with equal amounts of a DNA spike-in standard containing known barcodes. The barcodes are sequenced to quantify the ability of each F variant to infect cells in each antibody concentration, with barcode sequencing counts converted to fraction infectivity by normalizing them to the counts for the spike-in standard (see Methods). Details including all data for experimental replicates are shown here https://github.com/dms-vep/RSV_Long_F_DMS. **(b)** Correlation between effects of mutations on neutralization IC50 of subtype A (Long strain) or subtype B (B1 strain) F pseudoviruses in validation neutralization assays with nirsevimab IgG and Fab. Dashed gray line shows 1:1. **(c)** Correlation between effects of mutations in the deep mutational scanning of subtype A F with nirsevimab Fab or IgG versus effects on IC50 in validation neutralization assays using a subtype B (B1 strain) F. As predicted by the biophysical model, deep mutational scanning with the Fab is more predictive of effects of mutations in subtype B. Horizontal and vertical dashed lines indicate limits of detection. **(d)** Differences between subtype A and B at F sites 67 and 209 contribute to reduced potency of nirsevimab against subtype B. These neutralization curves are duplicated from Figure 3, and the same curves for the unmutated subtype A (Long strain) and subtype B (B1 strain) are plotted in both the top and bottom rows.

**Extended Data Figure 5.**
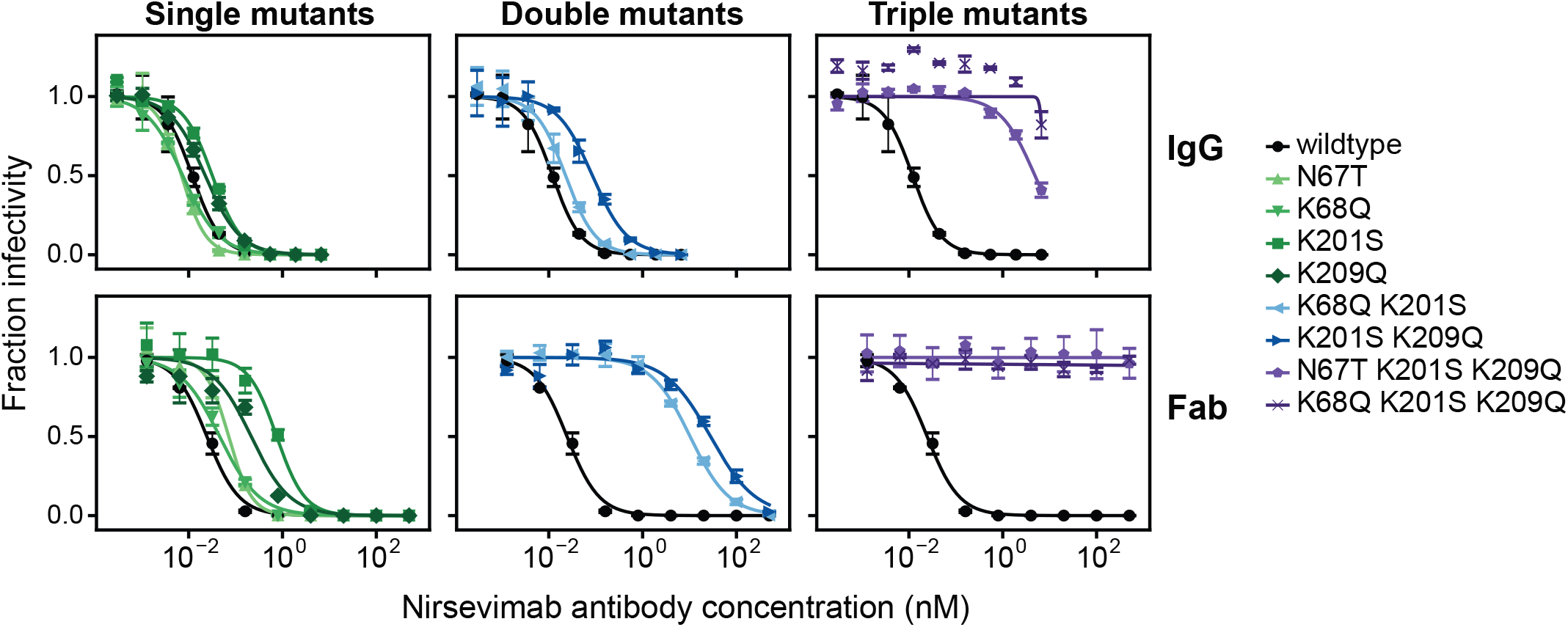
Mutations that individually reduce nirsevimab Fab but not IgG neutralization combine to also reduce IgG neutralization. Neutralization by nirsevimab IgG (top row) or Fab (bottom row) against pseudoviruses expressing RSV F from the subtype A Long strain with the indicated single, double, or triple mutations. Note that the same neutralization curve for the wildtype F from the Long strain against IgG or Fab is re-plotted in all three columns in each row to enable visual comparison of the mutants against wildtype.

**Extended Data Figure 6.**
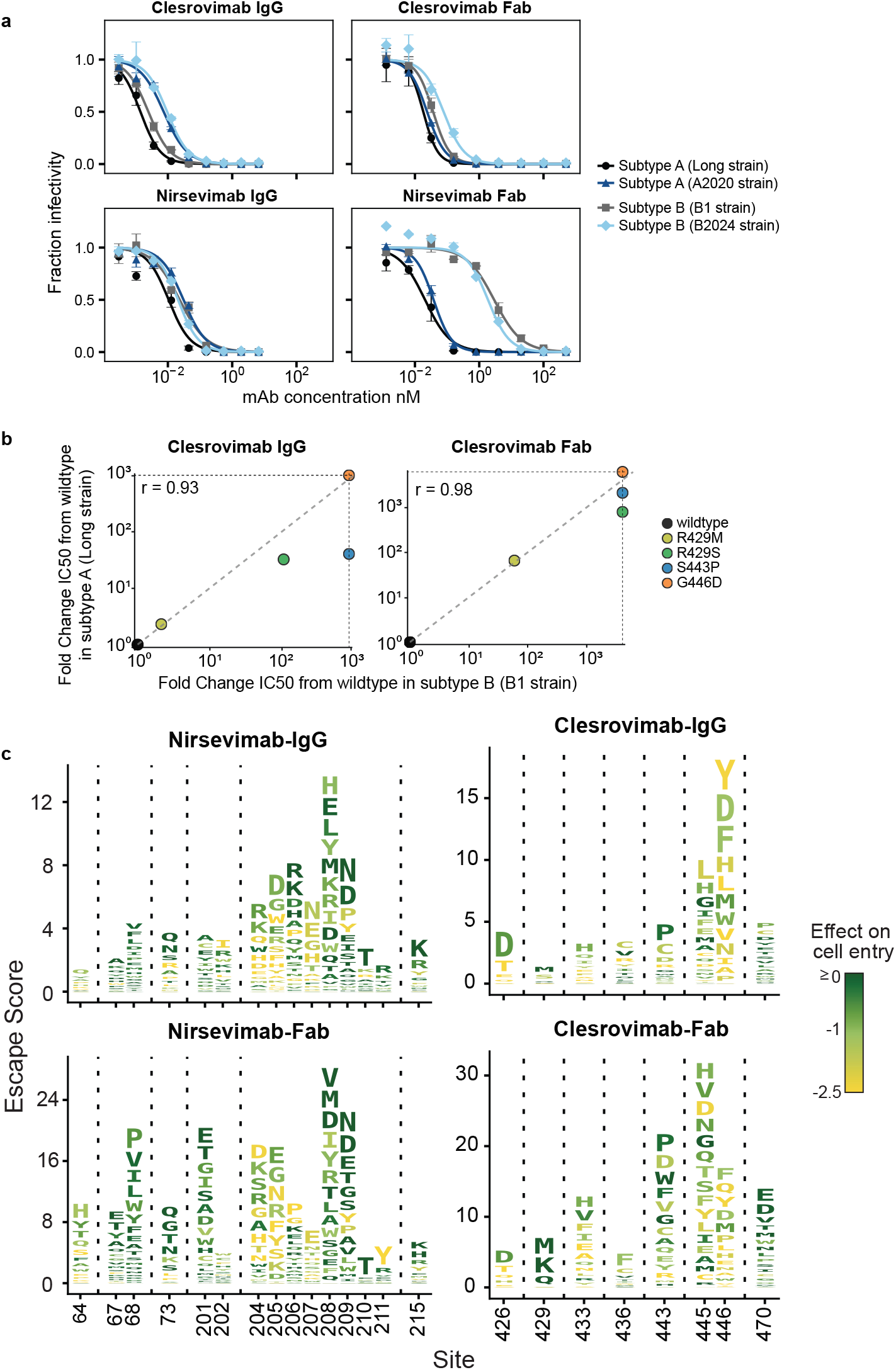
Mutations that affect clesrovimab neutralization have similar effects in subtypes A and B. **(a)** Neutralization curves for both the IgG and Fab forms of clesrovimab and nirsevimab IgG and Fab against pseudoviruses with F from two subtype A strains (the lab-adapted Long strain and the recent clinical strain A2020) and two subtype B strains (the lab-adapted B1 strain and the recent clinical B2024 strain). Points indicate the mean ± standard error of two technical replicates. **(b)** Correlation between effects of mutations on neutralization IC50 of subtype A (Long strain) or subtype B (B1 strain) F pseudoviruses in validation neutralization assays with clesrovimab IgG and Fab. Dashed gray line shows 1:1. **(c)** Mutations that reduce clesrovimab neutralization tend to be more deleterious for F’s cell entry function than mutations that reduce nirsevimab neutralization. Logo plots showing the effects of mutations at key sites on neutralization by nirsevimab or clesrovimab, measured by deep mutational scanning using IgG or Fab. The height of each letter is proportional to the reduction in neutralization a mutation to that amino acid causes in the deep mutational scanning. Mutations are colored by their effect on F-mediated cell entry in the absence of antibody as measured by deep mutational scanning, with dark green indicating no effect on cell entry and yellow indicating reduced cell entry. Therefore, yellow letters represent mutations that are likely to be deleterious to viral fitness even if they reduce antibody neutralization. Note that it is impossible to measure the effect on neutralization of mutations that fully ablate cell entry, so these plots only show mutations that retain at least some moderate level of cell entry (a cell entry effect > −2.5 in our measurements). See Figure 2 and associated hyperlinks to interactive plots for data on how all mutations at sites in the nirsevimab and clesrovimab epitopes affect cell entry function.

**Extended Data Figure 7.**
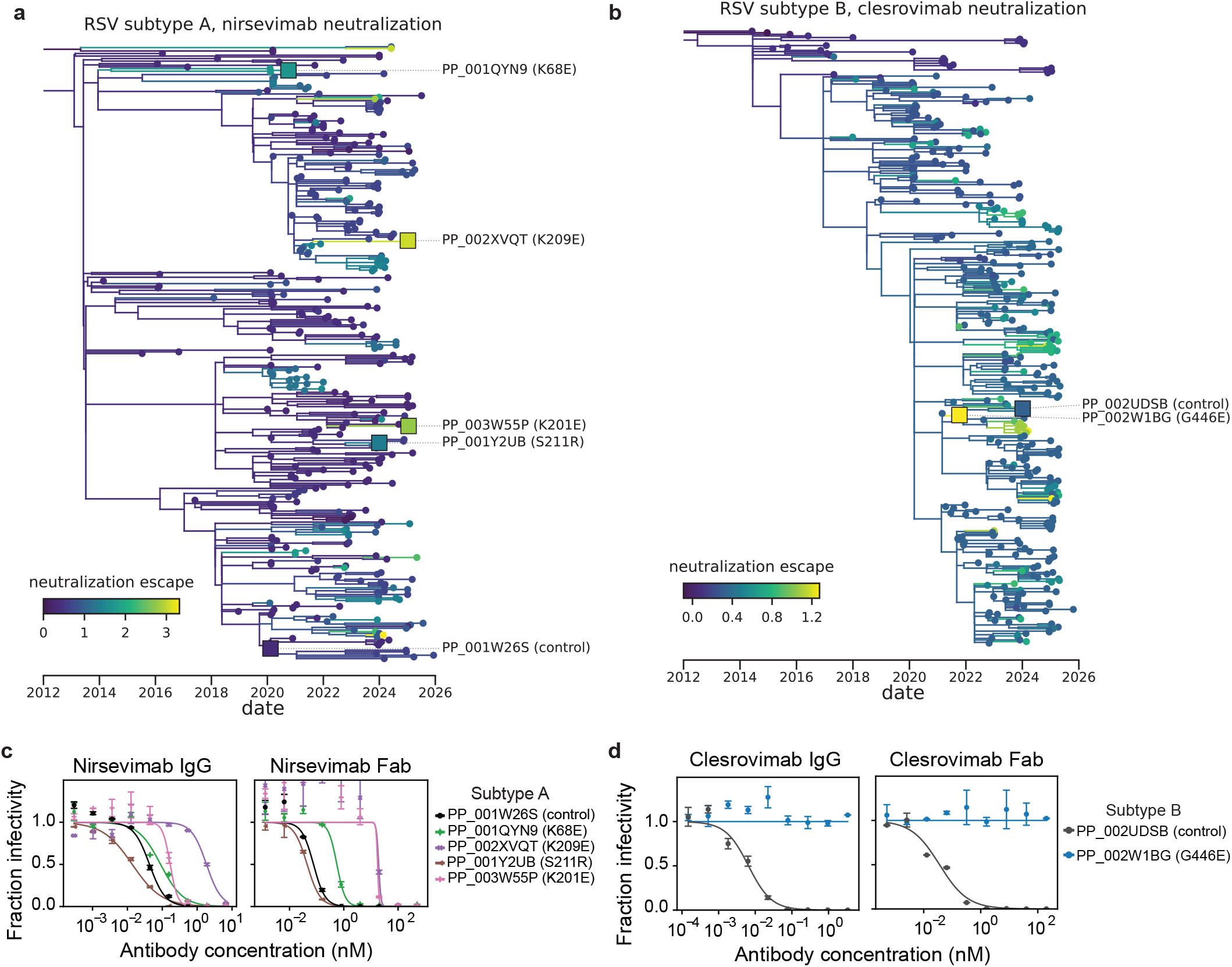
Additional phylogenetic trees and validation assays showing sporadic nirsevimab and clesrovimab resistance mutations in natural RSV sequences. This figure is like Figure 5 except it shows **(a)** trees of subtype A RSV F with nirsevimab escape scores, **(b)** trees of subtype B RSV F with clesrovimab escape scores, **(c)** neutralization assays validating reduced neutralization of natural subtype A strains by nirsevimab, and **(d)** neutralization assays validating reduced neutralization of a natural subtype B strain by clesrovimab. Note that subtype A strain PP_001Y2UB strain does not have reduced nirsevimab neutralization despite a high escape score driven by the S211R resistance mutation (which Figure 3d validates reduces nirsevimab neutralization in another subtype A strain); we speculate that neutralization of strain PP_001Y2UB might be affected by an epistatic interaction between S211R and a second R213S mutation it contains at a nearby site.

**Extended Data Figure 8.**
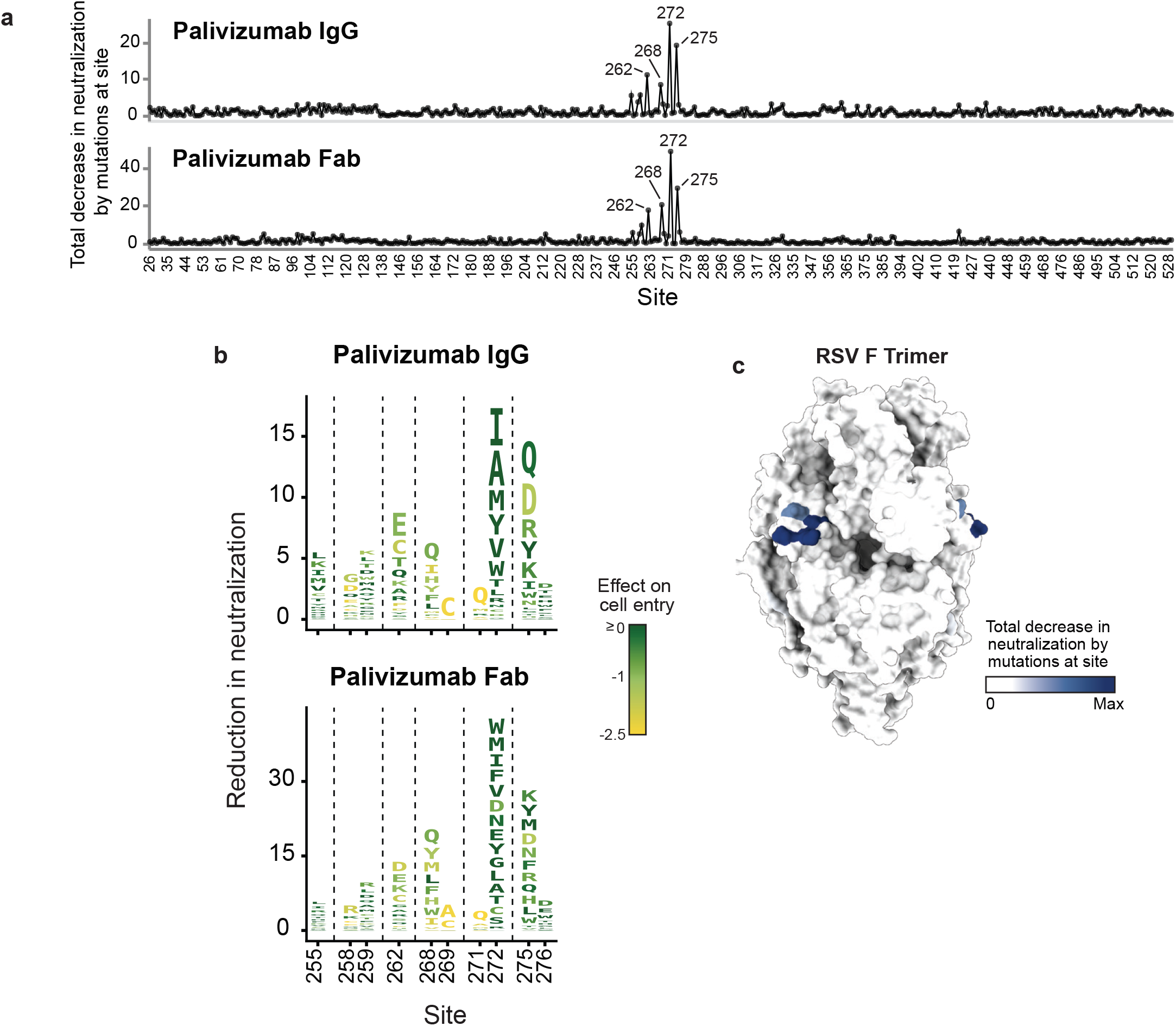
Effects of F mutations on palivizumab neutralization. **(a)** Total decrease in neutralization from all mutations at each site for palivizumab IgG or Fab as measured by deep mutational scanning. See https://dms-vep.org/RSV_Long_F_DMS/palivizumab_neutralization.html for interactive plots that show the effects of all mutations. **(b)** Logo plots showing the effects of mutations at key sites on neutralization by palivizumab measured by deep mutational scanning using IgG or Fab. The height of each letter is proportional to the reduction in neutralization a mutation to that amino acid causes in the deep mutational scanning. Mutations are colored by their effect on F-mediated cell entry in the absence of antibody as measured by deep mutational scanning, with dark green indicating no effect on cell entry and yellow indicating reduced cell entry. Therefore, yellow letters represent mutations that are likely to be deleterious to viral fitness even if they reduce antibody neutralization. It is impossible to measure the effect on neutralization of mutations that fully ablate cell entry, so these plots only show mutations that retain at least some moderate level of cell entry (a cell entry effect > −2.5 in our measurements). **(c)** Pre-fusion RSV F trimer (PDB 5UDC^13^) colored by the total effect of mutations at each site on neutralization by palivizumab Fab.

**Extended Data Figure 9.**
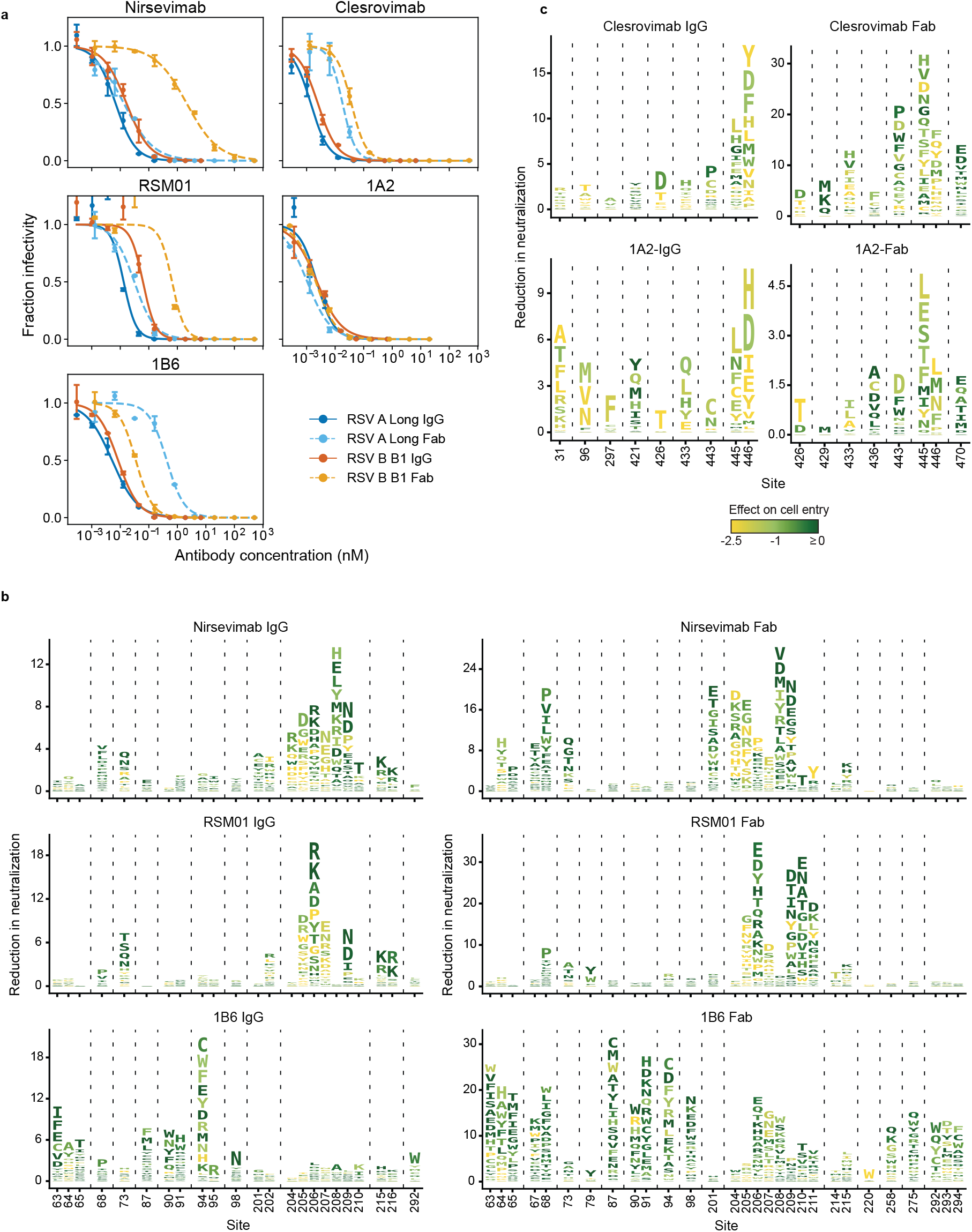
Comparison of mutational effects on neutralization by current (nirsevimab, clesrovimab) or candidate future (RSM01, 1A2, 1B6) clinical antibodies. **(a)** Neutralization curves for both the IgG and Fab forms of nirsevimab, RSM01, 1B6, clesrovimab, and 1A2 against pseudoviruses with F from the subtype A Long strain or subtype B B1 strain. Data for clesrovimab IgG and Fab are re-plotted from Extended Data Figure 6a to facilitate comparison to other antibodies plotted here. **(b)** Logo plots showing the effects of mutations at key sites on neutralization by apex-targeting antibodies (nirsevimab, RSM01 and 1B6), or **(c)** lateral face-targeting antibodies (clesrovimab and 1A2) as measured by deep mutational scanning using IgG or Fab. The height of each letter is proportional to the reduction in neutralization a mutation to that amino acid causes in the deep mutational scanning. Mutations are colored by their effect on F-mediated cell entry in the absence of antibody as measured by deep mutational scanning, with dark green indicating no effect on cell entry and yellow indicating reduced cell entry. Therefore, yellow letters represent mutations that are likely to be deleterious to viral fitness even if they reduce antibody neutralization. It is impossible to measure the effect on neutralization of mutations that fully ablate cell entry, so these plots only show mutations that retain at least some moderate level of cell entry (a cell entry effect > −2.5 in our measurements).

## Methods

### Data availability

All data and interactive figures are publicly available:

- Homepage for exploring interactive visualizations of deep mutational scanning data: https://dms-vep.org/RSV_Long_F_DMS/
- GitHub repository of all code used to analyze and visualize deep mutational scanning data, as well as processed data giving all sequencing counts and inferred mutation effects: https://github.com/dms-vep/RSV_Long_F_DMS
- Final measured effects of mutations on cell entry and anti-body neutralization after data QC: Supplementary File 1 and https://github.com/dms-vep/RSV_Long_F_DMS/blob/main/results/summaries/all_antibodies.csv
- Viral genome sequences and associated metadata were obtained from Pathoplexus and can be found at https://doi.org/10.62599/PP_SS_628.1, https://doi.org/10.62599/PP_SS_661.1, and https://doi.org/10.62599/PP_SS_662.1
- Previously published structures used for data visualization: PDB 5UDC^13^, PDB 6OUS^14^, PDB 9LLY^52^

### Code availability

A GitHub repository with all code used to analyze and visualize deep mutational scanning data, as well as processed data giving all sequencing counts and inferred mutation effects is available at https://github.com/dms-vep/RSV_Long_F_DMS and archived on Zenodo at DOI 10.5281/zenodo.21049419.

### Biosafety and biosecurity

We performed all experiments with pseudotyped lentiviral particles at biosafety level 2^60^. These pseudoviruses do not encode any viral proteins other than RSV F and therefore can only undergo a single round of cell entry and so are not fully replicative infectious agents capable of causing disease. The other viral proteins needed for the formation of pseudotyped lentiviral particles (RSV G, and the lentiviral Gag/Pol, Tat, and Rev) were provided during pseudovirus production by transfection of four separate helper plasmids and are not encoded in the viral genome. Therefore, this study did not generate any mutants of fully replicative biological agents capable of causing disease.

Our study quantifies how all single mutations to F affect neutralization by antibodies, including those in clinical use. However, RSV is already evolving under widespread pressure from these antibodies in the human population, with resistant strains regularly identified in breakthrough infections^3,4,6^. In the past, escape mutations have been identified from these breakthrough infections or by passaging authentic RSV virus in the presence of antibodies^3–6,14,22,26^. Our experiments systematically measure the effects of mutations outside the context of pathogenic virus, and therefore enable informed surveillance of how already ongoing evolution impacts clinical antibodies, as well as informing the design of new antibodies more resilient to viral resistance.

### Antibodies

The RSV monoclonal antibodies nirsevimab^13,19^, clesrovimab^14,40^, palivizumab^47^, suptavumab^2^, RSM01^11^, 1A2^52^ and 1B6^52^ were produced by Genscript as human IgG1 kappa isotypes and Fabs. Sequences were obtained from the referenced publications, structures in the Protein Data Bank or original patents. Palivizumab originated from patent US6955717B2. See https://github.com/dms-vep/RSV_Long_F_DMS/tree/main/supplemental_data/antibody-sequences for antibody amino acid sequences. The sequence of RSM01 was shared by the Gates Medical Research Institute.

### Plasmids and primers

All plasmid and primer sequences used in this study can be found at https://github.com/dms-vep/RSV_Long_F_DMS/tree/main/supplemental_data. All primers were ordered from Integrated DNA Technologies.

### Cell line handling

All cell lines (293T, 293T-TIM1^31^ and Takara Lenti-X™ 293T) were cultured in D10 media (Dulbecco’s Modified Eagle Medium supplemented with 10% heat-inactivated fetal bovine serum, 2 mM l-glutamine, 100 U/mL penicillin, and 100 mg/mL streptomycin) and cultured at 37ºC with 5% CO2.

### Design of deep mutational scanning libraries of RSV F

We created pseudovirus libraries containing nearly all possible single amino acid mutations to the ectodomain of the RSV F protein. We used unmutated parental F from the Long strain of RSV, which is a subtype A, lab-adapted strain isolated in the 1950s^32^. We codon optimized this sequence using the GenSmart codon optimization tool offered by GenScript and removed four amino acids from the cytoplasmic tail to increase pseudovirus titers^31^. We used a lentiviral backbone that contains an extended Gag sequence to enhance packaging of lentiviral genomes as previously described^61,62^. The plasmid map for the lentiviral backbone with codon-optimized RSV F sequence is at: https://github.com/dms-vep/RSV_Long_F_DMS/blob/main/supplemental_data/plasmids/4821_v5lp_phru3_forind-extgag_RSV_Long_F_GS4Opt_4aaCTdel.gb.

We aimed to include all single amino acid mutations in the RSV F ectodomain (residues 26-529; 504 x 19 = 9576 mutations). We also included 30 stop codons located at alternating positions from the start of the ectodomain as a negative control for cell entry measurements. We ordered a site-saturation variant library with these criteria from Twist Biosciences. The final Twist quality control report for the library is at: https://github.com/dms-vep/RSV_Long_F_DMS/blob/main/supplemental_data/Final_QC_Report_Q-392996_VariantProportion.csv.

### Cloning of deep mutational scanning plasmid libraries of RSV F

The RSV F library was designed to have all mutations to the ectodomain of F. However, 268 mutations were missing from the library produced by Twist. We added the missing mutations and also over-represented mutations in the nirsevimab binding site (amino acid residues 62-69 and 196-212) because we wanted to ensure inclusion and measurement of effects of all possible mutations in the epitope. We aimed to clone a “spike-in” plasmid library that contains these missing mutations and over-representation of mutations in the nirsevimab binding site using a mutagenesis PCR protocol^63–65^. We designed NNS primers for the nirsevimabtargeting sites with https://github.com/jbloomlab/CodonTilingPrimers and primers for missing mutations with https://github.com/jbloomlab/TargetedTilingPrimers. Forward and reverse primer pools were created by combining either forward or reverse NNS and targeted mutation primers at an equal molar ratio per codon for a final concentration of 5µM. Linear template of RSV F was made by digesting the lentiviral backbone with codon-optimized RSV F sequence with NotI-HF and NdeI. PCR mutagenesis was then performed as described previously^30^ with the only difference being that 7 PCR cycles were used for the mutagenesis PCR to reduce the resulting number of multi-mutants. After the PCR, the product was digested using DpnI for 20 minutes at 37°C to remove any leftover template. The PCR mutagenesis to generate the “spike-in” was performed in duplicate, once for library A and once for library B.

We then barcoded the library pools made by Twist Biosciences independently in two separate reactions to make library A and B respectively. These two biological replicates were handled separately for all subsequent experimental steps. The barcoding was performed as in Dadonaite et al.^30^ using primers containing a random 16 nucleotide sequence downstream of the RSV F stop codon. The only difference from Dadonaite et al. is that only 5 ng of template was used. The two pools of mutagenized RSV F for the “spike-in” were also barcoded separately. In all there were four separate barcoding PCR reactions. The lentiviral backbone (https://github.com/dms-vep/RSV_Long_F_DMS/blob/main/supplemental_data/plasmids/4016_V5LP_pHrU3_ForInd-Extgag_mcherry.gb)^62^ was digested with MluI-HF and XbaI at 37°C for 45 minutes then gel purified and purified using AMPure XP beads (Beckman Coulter Cat #A63881). The barcoded libraries were cloned into the lentiviral backbone at a 1:2 insert to vector ratio in a HiFi assembly for 1 hour at 50°C. The HiFi product was purified with AMPure XP beads and eluted in molecular grade water. The purified products were transformed into 10-beta electrocompetent cells (New England Biolabs Cat #C3020K) using a BioRad MicroPulser Electroporator, shocking at 2 kV for 5 milliseconds. 10 reactions of electroporation were performed per barcoded Twist library and 2 reactions per mutagenized spike-in library. Following the 1 hour recovery at 37°C, transformed cells were spun out of SOC and pooled and cultured in 150 mL LB medium for each Twist library and 100 mL LB medium for each spike-in library with ampicillin overnight at 37°C in a shaking incubator. Plasmids were extracted using the QIAGEN HiSpeed Plasmid Maxi Kit (QIAGEN Cat #12662).

Corresponding replicates of the Twist plasmid libraries and spike-in plasmid libraries were combined at a 1:1.25 Twist to spike-in molar ratio per codon because long-read PacBio sequencing of the plasmid libraries revealed this ratio results in the most even distribution of mutants in the combined libraries.

### Production of cell-stored deep mutational scanning libraries

Cells storing the deep mutational scanning libraries as single integrated RSV F containing genomes per cell were produced as described in prior work^30^ with a few changes (Extended Data Figure 1b). VSV-G pseudotyped lentiviruses were produced by transfecting four 10 cm dishes of 293T cells with the lentiviral backbone containing the barcoded RSV F libraries per library (5 μg per dish), lentiviral Gag/Pol, Tat, and Rev helper plasmids (1.25 ug per dish, AddGene: HDM-tat1b product ID 204154, pRC-CMV-Rev1b product ID 20413, HDM-Hgpm2 product ID 204152)^30^ and a VSV-G expression plasmid (1.25 ug per dish, AddGene pMD2.G product ID 12259) using BioT (Bioland Scientific, Cat #B01-02).

While previous deep mutational scanning studies have integrated into a specific clone of 293T-rtTA cells that were previously found to yield good pseudovirus titers for other viral entry proteins^30^, we found that this clone was unable to generate high titer RSV pseudovirus. We tested multiple clones of 293T-rtTA cells but found that the highest titer RSV pseudovirus produced from singly integrated cells was produced from Takara’s Lenti-X™ 293T Cell Line (Cat #632180). This clonal 293T cell line does not overexpress rtTA. Our lentiviral backbone includes a doxycycline-inducible promoter; in other deep mutational scanning studies which use 293T-rtTA cells and a similar backbone, doxycycline is added to induce expression from the lentiviral backbone^30^. We found that for production from RSV F cell stored libraries, pseudovirus titers were sufficiently high when expression was induced only from Tat at transfection. We found no additional benefit with transfection of rtTA and addition of doxycycline when producing pseudovirus. Therefore, we used Takara’s Lenti-X™ 293T Cell Line for the cell stored library. To make the cell stored library, the VSV-G pseudotyped viruses were used to infect Takara’s Lenti-X™ 293T Cell Line at an infection rate of <1% so that most transduced cells would receive only a single integrated genome. Transduced cells were selected using puromycin so that the final population of cells contained an integrated genome encoding a single barcoded variant of RSV F. These cells were expanded and frozen with at least 2e7 cells per aliquot and stored in liquid nitrogen for later use.

### Rescue of F and VSV-G expressing pseudovirus libraries

To rescue F expressing pseudoviruses from the integrated cells, 100 million cells were plated in 5-layer flasks. The following day, each flask was transfected with 118.75 ug of each helper plasmid (AddGene: HDM-tat1b product ID 204154, pRC-CMV-Rev1b product ID 20413, HDM-Hgpm2 product ID 204152) and 18.75 ug of RSV G expression plasmid (AddGene: HDM_RSV_Long_G_31AACTdel Product ID 237350). Xfect transfection reagent (Cat #631318) was used according to the manufacturer’s instructions (112.5uL Xfect transfection reagent and 7.5 mL buffer per 5-layer flask). At 48 hours post transfection, the supernatant was filtered through a 0.45um SFCA Nalgene 500 mL Rapid-Flow filter unit (Cat #09-740-44B). Filtered supernatant was then concentrated by ultracentrifugation with a 20% sucrose cushion in HBSS (Fisher Cat# 14025092) at 100,000 g for 1 h at >20ºC. The pseudoviruses pelleted to the bottom of the cushion, after disposal of the supernatant, pellets were resuspended in D10. Aliquots of concentrated F-expressing pseudoviruses were flash frozen as previously described^31^ and stored at −80°C for use in downstream selection experiments.

To rescue VSV-G expressing pseudoviruses from integrated cells, 100 million cells were plated in 5-layer flasks. The following day, each flask was transfected with 37.5 ug each helper (AddGene: HDM-tat1b product ID 204154, pRC-CMV-Rev1b product ID 20413, HDM-Hgpm2 product ID 204152) and VSV-G expression plasmid (AddGene pMD2.G product ID 12259). BioT transfection reagent was used according to the manufacturer’s instructions. At 48 hours post-transfection, the supernatant was filtered through a 0.45um SFCA Nalgene 500 mL Rapid-Flow filter unit and concentrated using LentiX concentrator (Takara, Cat #631232) at a 1:3 virus to concentrator ratio, incubating at 4°C for 3 hours and spinning at 1500 x g and 4°C for 45 minutes. Following centrifugation, supernatant was discarded and viral pellets resuspended in D10. Aliquots of concentrated VSV-G expressing pseudoviruses were frozen at −80°C for later use.

### Long-read PacBio sequencing for variant-barcode linkage

To link the barcode sequences with the mutations found in RSV F we used long-read PacBio sequencing of the lentiviral genomes in pseudoviruses made from the cell stored libraries. We performed PacBio barcode-mutation linking after generating the singly integrated cell libraries because template switching of the pseudodiploid lentiviral genome during reverse transcription can alter barcode-variant pairings relative to the original plasmid pool. Sequencing the integrated library ensures that barcode-variant linkages reflect those present in the actual integrated provirus. This process has been described previously^30^. For this library we made a few alterations. 1×10^6^ 293T-TIM1 cells were plated in each well of 6-well plates coated with poly-L-lysine. The next day, 30 million TU’s of VSV-G expressing library pseudoviruses were used to infect cells (6 wells for each library at 5 million TU/well). At 12 hours post-infection, the non-integrated reverse-transcribed lentiviral genomes were recovered by miniprepping the 293T-TIM1 cells as described in prior work^30^. Amplicons for long-read sequencing of the miniprepped genomes were prepared by following a previously described approach^30,65^. The PCR reactions for each library were combined and amplicon length was verified by TapeStation prior to sequencing. Libraries were each sequenced on a single SMRT cell with a movie length of 30 hours on a PacBio Sequel IIe sequencer. To maximize identification of all variants present, each library was sequenced a second time.

We used the *dms-vep-pipeline-3* package (https://github.com/dms-vep/dms-vep-pipeline-3), to process sequencing data. To link specific mutations with each barcode, PacBio circular consensus sequences were aligned to the unmutated RSV F reference sequence using the *Alignparse* package^66^. Reads were filtered out if they aligned poorly, had a higher than expected number of mutations in the unmutated regions, did not contain a barcode or were the result of strand exchange. Consensus sequences for each barcode/variant sequence were constructed using *alignparse*, while requiring a minimum of three CCS reads and a max cutoff of 0.2 for any minor variants within the consensus. The final bar-code/variant lookup tables were used as a reference for all downstream analyses of short-read Illumina sequencing of the barcodes only.

The final barcode-variant table can be found here: https://github.com/dms-vep/RSV_Long_F_DMS/blob/main/results/variants/codon_variants.csv

For full details on the analysis, see these notebooks for:

- Analyzing the PacBio CCS’s: https://dms-vep.org/RSV_Long_F_DMS/notebooks/analyze_pacbio_ccs.html
- Building PacBio consensus sequences: https://dms-vep.org/RSV_Long_F_DMS/notebooks/build_pacbio_consensus.html
- Building the final barcode-variant table: https://dms-vep.org/RSV_Long_F_DMS/notebooks/build_codon_variants.html

The final libraries contained 57,888 and 64,078 unique barcoded variants for LibA and LibB, respectively that covered 99 and 99.4% of all amino acid mutations (Extended Data Figure 1c). More than half of the variants contain a single amino acid mutation, with the others having zero or multiple mutations (Extended Data Figure 1d).

### Measuring effects of F mutations on cell entry

To measure effects of F mutations on cell entry, we generally followed the approach described in Dadonaite et al.^30^ with the following modifications. We infected 293T-TIM1 cells with the pseudovirus libraries displaying the RSV F protein mutants alongside parallel infection of 293T-TIM1 cells with control pseudovirus displaying VSV-G to make their cell-entry independent of RSV F protein function (Extended Data Figure 2a). 2×10^6^ 293T-TIM1 cells were plated in each well of 6-well plates coated with poly-L-lysine. The next day, we infected cells with ~12×10^6^ TU total (3×10^6^ TU per well) of F pseudovirus library or ~3×10^7^ TU total (5×10^6^ TU per well) of VSV-G pseudovirus library. After addition of pseudovirus, cells infected with the F pseudovirus library were spun at 900g for 3 hours at 30C. For some selections, 20ug/mL DEAE-dextran was added to the media at the time of infection as this increases pseudovirus titers. However, poor cell health was found to contribute to reduced recovery of infecting barcodes, so DEAE was not used for follow-up selections. At 12 hours post infection, the non-integrated reverse-transcribed lentiviral genomes were recovered by miniprepping the cells. To prepare the amplicons for Illumina sequencing with dual indexing, PCR was performed as described in Yu et al^65^. Samples were pooled in equal DNA amounts and run on a 1% agarose gel. The correct size band was excised, purified with Ampure XP beads, diluted to a concentration of 5 nM, and sequenced on an Illumina NextSeq 2000 (with P3 reagent kit) or NovaSeq X Plus system.

We quantified the effects of mutations as the log2 cell entry of each mutant relative to the unmutated F, using global epistasis models^33,34^ to jointly analyze the single and multi-mutant data. Briefly, Illumina sequencing reads were aligned to the barcode variant table generated from the PacBio CCS described above. We next compared the frequency of barcodes between the VSV-G and F_mutant_ selections using the package *dms_variants* (https://github.com/jbloomlab/dms_variants) as previously described^30^. Cell entry scores for each variant were calculated using log enrichment ratio: *log*_2_ ([*n*^*v*^_*post*_ / *n*^*wt*^_*post*_]/[*n*^*v*^_*pre*_ / *n*^*wt*^_*pre*_]), where *n*^*v*^_*post*_ is the count of variant *v* in the F-pseudotyped infection (postselection condition), *n*^*v*^_*pre*_ is the count of variant *v* in the VSV-G-pseudotyped infection (preselection condition), and *n*^*wt*^_*post*_ and *n*^*wt*^_*pre*_ are the counts for wild-type variants. Positive cell entry scores indicate that a variant is better at entering the cells compared to the unmutated parental F, and negative scores indicate entry worse than the parental F. As expected, variants with only synonymous mutations had wildtype-like cell entry scores of zero, variants with stop codon mutations had highly negative scores, and variants with amino acid mutations had scores ranging from wildtype-like to highly negative (Extended Data Figure 2b). To calculate the mutation-level cell entry effects, a sigmoid global-epistasis function was fitted to variant entry scores after truncating the values at a lower bound of the median functional score of all variants with stop codons, using the *multi-dms* software package. For the mutation effects, values of zero mean no effect on cell entry, negative values mean impaired cell entry, and positive values mean improved cell entry. To generate the final cell entry effect values for each mutation, we performed a total of three technical replicates for each library, for a total of six functional selections (three each from LibA and LibB). The effects for each mutation were then averaged from the six functional selections. To filter out low quality or noisy data, we required each mutation to occur with two unique barcodes and removed any mutation that had a high standard deviation between replicates. The measured cell entry effects were highly correlated both between replicates and libraries (Extended Data Figure 2c,d). The final cell entry effect values reported in the figures correspond to the average effect across libraries and replicates.

### Measuring effects of F mutations on antibody neutralization

To measure effects of F mutations on antibody neutralization, we used a previously described method^30^ with a few modifications detailed here. 2×10^6^ 293T-TIM1 cells were plated in individual wells of poly-L-lysine coated 6-well plates. The next day, ~1.5-2×10^6^ TU of F pseudovirus library were incubated with D10 media (no antibody control) or antibody for one hour prior to adding to cells. For monoclonal antibodies, these incubations were performed in a total volume of 2 mL to reduce the possibility of ligand depletion given that these antibodies have very high potency. Antibody concentrations were selected that generally corresponded to a range at which 50% of variants were neutralized, up to 99.5%. During DNA template extraction, we spiked in DNA plasmid containing eight known barcodes that would correspond to ~1% of the reads in the no-antibody control. This DNA plasmid spike-in allowed us to estimate the amount of neutralization each antibody condition had relative to the no antibody control, as previously described (Extended Data Figure 4a)^30,65^.

Following Illumina sequencing of the barcodes, the data were analyzed as described previously. Briefly, the fraction infectivity retained at each antibody concentration was calculated from the barcode counts of the DNA standard. Then, the *polyclonal* (https://jbloomlab.github.io/polyclonal/)^67^ software was used to fit neutralization curves and estimates of mutation effects on neutralization. We filtered the effects of mutations on antibody neutralization to retain only mutations with at least two unique barcodes, excluded mutations with very low cell entry scores, and excluded mutations that had high standard deviations in their effect on neutralization across replicates. The reported effects of mutations on antibody neutralization are the average across all replicates (always at least two different experimental selections with each of the two independent libraries, LibA and LibB). For example notebooks showing these analyses, see https://dms-vep.org/RSV_Long_F_DMS/notebooks/fit_escape_antibody_escape_RSV-F-LibA-250402-Nir.html (for analysis of an individual selection experiment) and https://dms-vep.org/RSV_Long_F_DMS/notebooks/avg_escape_antibody_escape_Nirsevimab-IgG.html (for average effects across experiments for an antibody).

### Validation of cell entry effects using individual pseudoviruses

To validate the effects of mutations on cell entry, we generated a set of plasmids expressing RSV F within a lentiviral backbone that each contained different single F mutations. The parental RSV F amino acid sequence is identical to the unmutated RSV F sequence used in the lentiviral vector described above for deep mutational scanning. We selected mutations spanning a range of entry effects, which included N67I, S215V, S215P, S398L, E87N, and D486N. We transfected them into 293T cells, along with helper plasmids (AddGene: HDM-tat1b product ID 204154, pRC-CMV-Rev1b product ID 20413, HDM-Hgpm2 product ID 204152)^30^ and RSV G in an expression plasmid (AddGene: HDM_RSV_Long_G_31AACTdel

Product ID237350). 48 hours later, supernatants were filtered through 0.45uM filters to remove cell debris. The pseudoviruses were titrated on 293T-TIM1 cells using flow cytometry to measure TU/mL as previously described^31^. Average titers for two replicates of each pseudovirus were then compared to the average unmutated RSV F titer.

### Analysis of cell entry effects of amino acid differences in natural RSV sequences relative to the strain used for deep mutational scanning

An alignment of RSV F protein sequences for subtype A and B was generated on October 27, 2025 from the RSV Nextstrain workflow^45^. We identified amino acid differences relative to the lab-adapted subtype A Long strain used in the deep mutational scanning - for more details see https://github.com/dms-vep/RSV_Long_F_DMS/tree/main/notebooks/sequence_variation. This was used to compare the distribution of cell entry effects for mutations observed in natural sequences to the distribution of cell entry effects for all mutations measured by deep mutational scanning and calculate the percentage of natural sequences with each mutation as shown in Extended Data Figure 3.

### Measuring effects of single amino acid mutations on antibody neutralization

We chose previously known and newly identified resistance mutations that had a range of effects on antibody neutralization for nirsevimab and clesrovimab. These mutants were chosen to evaluate the impact of mutations in subtype A and B backgrounds on neutralization by antibody IgG and Fab and to validate the effects of mutations measured by deep mutational scanning on antibody neutralization. The single amino acid mutations were made in a subtype A background (Long strain, also used for deep mutational scanning) and a subtype B background (B1 strain) in an expression vector. The plasmids were generated by Twist Biosciences. Pseudoviruses were produced and neutralization was measured as previously described^31^. All F constructs included the full cytoplasmic tail and were paired with RSV G in an expression plasmid (AddGene: HDM_RSV_Long_G_31AACTdel Product ID 237350) for these transfections. All plasmid maps can be found here: https://github.com/dms-vep/RSV_Long_F_DMS/tree/main/supplemental_data/plasmids/.

Mutations impacting nirsevimab neutralization made in sub-type A included (N67T, K68N, K68Q, D73N, K201S, K201T, P205S, V207E, K209D, K209Q, Q210T, S211R, S215K) and subtype B (K68N, K68Q, D73N, N201S, N201T, P205S, Q209D, Q210T, S211R, S215K). V207E did not produce usable pseudovirus titers in the B1 background. These are shown in Figure 3 and Extended Data Figure 4.

Mutations T67N and Q209K were also made in subtype B, which are mutations to the amino acid residue found in the subtype A Long strain. Data for these mutants along with the corresponding mutations in subtype A (N67T and K209Q) are shown in Extended Data Figure 4. Subtype A K68Q and K201S and subtype B K68Q and N201S are shown in Figures 1, 3 and Extended Data Figure 4.

The Long and B1 wildtype neutralization curves shown in Figure 3d,f are representative - all replicates can be found here https://github.com/dms-vep/RSV_Long_F_DMS/tree/main/non-pipeline_analyses/validations. The correlation plots in Figures 3 and 4 show fold change IC50 from wildtype using a matched wild-type from the specific experimental date for the mutations and the wildtype point plotted in the correlation plots is a geometric mean IC50 of all wildtype replicates.

Mutations impacting clesrovimab made in subtype A and B included R429M, R429S, S443P and G446D and are shown in Figure 4 and Extended Data Figure 6. Pseudoviruses expressing F called “A2020” and “B2024” in Extended Data Figure 6a are F sequences from natural strains that are broadly representative of recent subtype A and B strains^31^, and have the GenBank accession numbers PP495954.1 and PP660445.1, respectively.

For all plotted neutralization curves, points represent the mean and standard error of at least two replicate measurements.

### Interactive Nextstrain phylogenetic trees with antibody-escape scores

We integrated computer code into the Nextstrain^45^ RSV view to enable scoring of natural RSV sequences for antibody resistance and visualization of the results on phylogenetic trees. A GitHub repository with the computer code implementing this scoring is at http://github.com/nextstrain/rsv. Briefly, the pre-existing Nextstrain view downloaded all RSV sequences from Pathoplexus^46^ (currently >60,000), and built separate subtype A and B phylogenetic trees subsampled to roughly ~3,000 sequences designed to be representative across time and countries. We added computer code that assigns each of these sequences an “escape score” calculated from the deep mutational scanning data as either the sum of the effect of all of its constituent mutations or the max effect of any of its mutations on neutralization by the Fab or IgG form of nirsevimab or clesrovimab. The interactive phylogenetic trees can then be colored by these escape scores using the dropdown “Color By” option on the left toolbar (eg, see https://nextstrain.org/rsv/b/F-antibody-escape/6y?c=Nirsevimab-Fab_total_escape); there is also an option to label sequences by their top escape mutation. We integrated this scoring into the Nextstrain RSV builds for the F sequences and the full genome; note that there are also builds that emphasize different timeframes: all-time, the last 6 years, and the last 3 years. The builds are updated to include the latest available sequences, so going to views linked above will show the latest sequence data.

We also created new builds called “F-antibody-escape” in the Nextstrain RSV views that show trees subsampled to include all sequences with high nirsevimab or clesrovimab escape scores. Unlike the “F” builds, these “F-antibody-escape” builds over-represent the frequency of resistant strains. The “F-antibody-escape” trees do not provide an accurate view of the prevalence of resistance mutations; however, when you want to identify all of the top resistant strains (which otherwise could be dropped during subsampling) then these builds should be preferred. Which build is displayed can be selected using the “change dataset” option on the left toolbar.

### Validation of strains with predicted neutralization resistance

A subset of natural RSV F sequences with predicted resistance to nirsevimab or clesrovimab neutralization identified using the ‘Interactive Nextstrain phylogenetic trees with antibody-escape scores’ were codon optimized and subsequently cloned into an expression vector by Twist Biosciences. F sequences with predicted resistance to nirsevimab Fab or IgG included (PP_002XVQT, PP_002KSRJ, PP_002QMLP, PP_002WHEU, PP_002WWH8, PP_002SUFP, PP_001QYN9, PP_001Y2UB, PP_003W55P). Sequences with predicted resistance to clesrovimab Fab or IgG included (PP_002W1BG, PP_001WGC0, PP_001Y62S, PP_001ZQ7W). PP_001W26S and PP_002UDSB correspond to subtype A and B controls also referred to in our prior study as A2020 and B2024, respectively^31^. A complete list of these key sequences is available in Pathoplexus under SeqSet PP_SS_628.1^68^, and all sequences displayed in the trees in Figure 5 and Extended Data Figure 7 are available in Pathoplexus under SeqSet PP_SS_661.1^69^ for subtype A and SeqSet PP_SS_662.1^70^ for subtype B. All plasmid maps can be found at: https://github.com/dms-vep/RSV_Long_F_DMS/tree/main/supplemental_data/plasmids/strain%20validations. These were used to generate RSV pseudoviruses expressing the F from the natural sequences paired with RSV G expression plasmid (AddGene: HDM_RSV_Long_G_31AACTdel Product ID237350) as previously described^31^. Producing RSV pseudoviruses with F from natural sequences paired with all the same G ensures we do not introduce any variability from G. These pseudoviruses were then used in neutralization assays with nirsevimab or clesrovimab IgG and Fab^31^. To ensure reliable neutralization curves, we established a cut-off of 400,000 RLU/well for the no antibody, virus-only control. For strains with titers near the cut-off, extra controls were included to ensure escape was specific to the monoclonal antibody and not attributable to experimental artifacts.

### Structural analysis

UCSF ChimeraX^71^ was used for structural visualizations. All Protein Data bank accession IDs used are included in figure legends.

## Supplementary Files

### Supplementary File 1. Effects of mutations on cell entry and monoclonal antibody neutralization

This CSV file is also available at https://github.com/dms-vep/RSV_Long_F_DMS/blob/main/results/summaries/all_antibodies.csv

### Supplementary Appendix: biophysical model of how bivalent binding shapes impact of viral mutations on IgG versus Fab neutralization

Here we describe a biophysical model of how the effect of a mutation to a viral antigen on neutralization by an antibody depends on the antibody’s affinity for the antigen. The two key effects modeled here are the bivalent binding of the IgG (but not Fab) and the ligand depletion (titration) that can put a floor on the lowest measurable neutralization value. The model is illustrated conceptually in Figure 1a, and is used to generate the simulated data in Figure 1b that is compared to the actual experimental data in Figure 1c.

See https://github.com/jbloomlab/IgG-vs-Fab-neutralization for the computer code implementing the model, and https://jbloomlab.github.io/IgG-vs-Fab-neutralization/notebook.html for an interactive notebook describing the model.

### Fab Binding

Let the binding affinity of the Fab have dissociation constant *K*_*D*_. There are two states, unbound (*U*) and bound (*B*). The Boltzmann weights for these two states as a function of the molar concentration *c* of Fab are *w*_*U*_(*c*) = 1 and 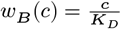. So the probability of the unbound state as a function of *c* is 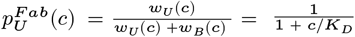. This two-state partition-function formulation for monovalent binding follows classic multivalency treatments^72,73^.

### IgG binding

For the IgG, we assume each arm of the bivalent IgG has the same monovalent affinity for the viral antigen as the Fab described above. However, once the first arm binds, the effective concentration *c*_*eff*_ of the second arm is set by the density/spacing of viral antigens and the reach of the antibody. Here we do not consider those factors (antigen density, antibody reach, etc) separately, but just collapse them into the single effective concentration of the second arm after the first has bound, *c*_*eff*_. The definition of an effective local concentration for the second arm, set by geometry/linker statistics rather than bulk concentration, mirrors prior work^28,74,75^. In realistic scenarios, if bivalent binding is possible then *c*_*eff*_ ≫ *c* since the effective concentration of the second Fab arm in proximity to the viral antigen after the first arm has bound is much higher than the overall IgG concentration (note that concentrations should be in *molar* units rather than ng/ml in order to allow the same concentration scale to be used for Fab and IgG, since they have different molecular weights).

There are now three possible states, unbound (*U*), a single Fab arm bound (*B*_1_), and both Fab arms bound (*B*_2_). The Boltzmann weights for these three states as a function of the molecular concentration of IgG are *w*_*U*_(*c*) = 1, 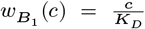, and 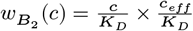. Further, note that state *B* has a multiplicity of two since either of the Fab arms can bind^73^. So the probability of the unbound state as a function of *c* is (accounting for the multiplicity of two for *B*_1_) 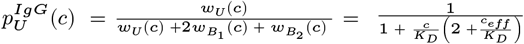. Note that the effective dissociation constant of the IgG is 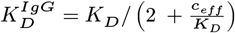.

### Assumption: binding affinity equals neutralization potency

In what follows, we assume that binding affinity equals neutralization potency, such that the neutralization IC50 of the Fab is equivalent to *K*_*D*_. In other words, we equate fraction infectivity with fraction of viral epitopes (*p*_*u*_); as described immediately below, deviations can occur under multi-hit models. Specifically, how fraction bound (binding affinity) relates to neutralization potency depends on how many antibodies must bind to a virion in order to neutralize it. The quantitative relationship between binding affinity and neutralization potency as well as its impact on the neutralization curve slope can be modeled under assumptions about the number of spikes per virion and the number that must be bound to neutralize^76–80^. Here we are going to *ignore* this issue for now, and just conflate neutralization IC50 with Fab *K*_*D*_. This is a quantitative simplification, but should not affect the main qualitative trends— and has the advantage of not requiring us to estimate difficult to determine quantities such as how many spikes there are per virion and what fraction of them must be bound for neutralization. Note that in the limiting case when there is just one spike per virion and binding one site on it is sufficient for neutralization, then ignoring this issue has no effect.

### Estimating *c*_*eff*_ from the IgG versus Fab IC50

We can estimate *c*_*eff*_ (the parameter that describes the extent of avidity from bivalent binding) from the IC50 (or midpoint on the neutralization curve) of the IgG versus Fab. In particular, let *mFab* and *mIgG* be the molar concentrations at the midpoint (IC50) of the Fab versus IgG neutralization curves, respectively. Then *K*_*D*_ = *mFab*and 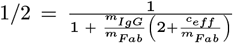 Solving for *c*_*eff*_ yields 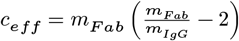.

### Impact of mutation(s)

Consider one or more mutations that cause a fold change in Fab IC50 of *f*_*mut*_. For instance, a mutation that increases the Fab IC50 by 20-fold would have *f*_*mut*_ = 20. Note that mutations that cause a fixed change in the free energy of binding are expected to cause the same fold change in Fab IC50 in any genetic background. In the presence of these mutation(s), the fraction of Fab not neutralized becomes 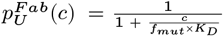 and the fraction of IgG not neutralized becomes 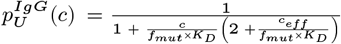.

### Complication: accounting for ligand depletion

Everything above is written in terms of the antibody (Fab or IgG) concentration *c*. In fact, *c* in these equations refers to the *free* concentration of antibody. But when the concentration of viral antigen is comparable to or exceeds the dissociation constant of the antibody (*K*_*D*_ for the Fab and 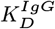 for the IgG), there can be “ligand depletion,” which is the term we use here for when a large fraction of the antibody is bound to the viral antigen. When this occurs, much of the antibody gets “soaked up” by binding to viral antigen, and so the actual concentration of free antibody in the neutralization assay represented by *c* in the above equations is appreciably less than the total concentration added to the reaction. This phenomenon is nicely described in detail by Jarmoskaite et al (2020)^27^, who refer to it as the “titration regime.”

To quantitatively deal with this, we need to define some additional variables. Let *c*_*total*_ be the total antibody concentration added to the assay volume, let *c*_*free*_ be the concentration of antibody that is not bound to viral antigen (free), and let *c*_*bound*_ be the concentration of antibody that is bound to viral antigen. Also, let *v*_*total*_ be the total concentration of viral antigen (quantified in terms of number of the relevant epitope per unit volume), let *v*_*free*_ be the concentration of free viral antigen (not bound by antibody), and note that concentration of bound viral antigen is *c*_*bound*_. We have the following equations: *c*_*total*_ = *c*_*free*_ + *c*_*bound*_ (mass balance for the antibody), *v*_*total*_ = *v*_*free*_ + *c*_*bound*_ (mass balance for the viral antigen), and 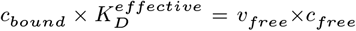 (there is an equilibrium with respect to the free antibody and antigen concentration). Note that for the Fab, 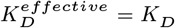 and for the IgG,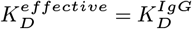.

Solving this system of three equations yields^27^ 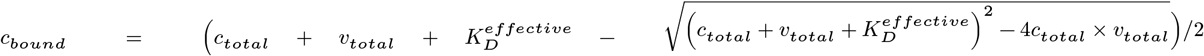 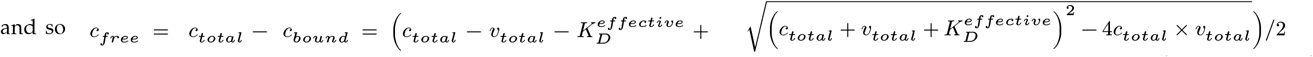.

Note that in the limit where the viral antigen concentration is limiting and so we are outside the ligand depletion regime (eg, 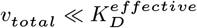), then we just have *c* _*free*_ ≈ *c* _*total*_. However, when the viral antigen concentration is comparable to or greater than 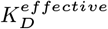, then it is important to substitute *c* _*free*_ for *c* in all of the equations above.

We can also calculate the actual observed IC50s accounting for ligand depletion. The IC50 (under our assumption above that this is when half of viral epitopes are bound) occurs when 1/2 = *c*_*bound*_ / *v*_*total*_. It turns out that this can be neatly solved to determine the actual observed IC50 (accounting for ligand depletion) is 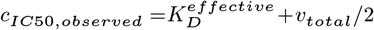, whereas the ideal IC50 if there is no ligand depletion (when 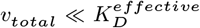), is just 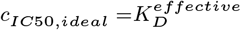. The floor on the measurable IC50 (eg, the lowest value that can be measured) is therefore *v*_*total*_/2.

### Realistic parameter estimates

We ballpark (these are rough estimates) realistic parameter values for nirsevimab and RSV F from our experiments (Figure 1c); the model-simulated data in Figure 1b use these parameters with the subtype A *K*_*D*_ estimates for the high-affinity panel and the subtype B *K*_*D*_ estimates for the low-affinity panel, and all other parameters shared between the panels as they are not expected to be subtype-specific. For the Fab IC50, which is represented by *m*_*Fab*_ or *K*_*D*_ in the above equations:

- subtype A (Long) strain: *K*_*D*_ ~ 0.01 nM
- subtype B (B1) strain: *K*_*D*_ ~ 1 nM

We also need to estimate *c*_*eff*_; however, doing that is complicated by the fact that at least for subtype A, the IgG neutralization assays are likely in the ligand depletion range (what Jarmoskaite et al^27^ call the “titration regime”) where the IgG IC50 (*m*_*IgG*_) cannot be measured accurately since the concentration of the viral antigen protein likely is comparable or greater than the true IgG IC50.

So we will use neutralization measurements against the subtype B (B1) strain as any IgG ligand depletion should be less here due to the lower potency, and we assume that the actual avidity (potential for bivalent binding) captured in *c*_*eff*_ should not be strain dependent. Against this strain, we have *m*_*IgG*_ ~ 0.01 nM and *m*_*Fab*_ ~ 1 nM. So this gives *c*_*eff*_ ~ 100 nM. However, there is probably still ligand depletion in the measurement of *m*_*IgG*_ since the subtype B IgG neutralization is just as good as subtype A, so we will add another factor of 10 and use *c*_*eff*_ ~ 1000 nM (note this remains a very rough estimate and real value could be higher).

The hardest parameter to estimate is the number of viral epitopes *v*_*total*_ which determines if there is ligand depletion. Roughly, we might estimate 10^6^ infectious particles per ml, with 10^2^ non-infectious particles for each infectious one, 10^2^ epitopes per particle. In that case, we have 10^13^ epitopes / liter, which gives *v*_*total*_ ~ 0.01 nM. Note that this estimate, which was used in Figure 1b, is very much a rough estimate and could vary by an order of magnitude or more in either direction.

### Interactive visualization of IgG and Fab neutralization and how it is affected by mutations

Go to https://jbloomlab.github.io/IgG-vs-Fab-neutralization/chart.html for an interactive chart initialized with the ballpark-realistic parameter values above for the subtype A strain of how nirsevimab neutralization differs between the Fab and IgG, and how this is impacted by mutations. In the interactive chart, you can use the sliders to adjust the Fab affinity (*K*_*D*_), the avidity (*c*_*eff*_), the effect of the viral mutation(s) (*f*_*mut*_), and the viral epitope concentration (*v*_*total*_). As you do so, you can see how the neutralization curves and IC50s change. Note for the IC50s, the interactive chart shows both the actual observed IC50 (accounting for ligand depletion) as well as what would be the ideal IC50 in the absence of any ligand depletion.

## Acknowledgements

We thank Richard Neher and James Hadfield for assistance with the Nextstrain visualizations. We thank Terry Stevens Ayers, Michael Boeckh and Alex Greninger for reviewing the manuscript.

## Author Contributions

Conceptualization: CALS, TEM, HYC, JDB; Methodology: CALS, TEM, LK, JDB; Investigation: CALS, TEM, GJ; Data curation: CALS, TEM, JDB; Visualization: CALS, TEM, JDB; Writing-Original Draft: CALS, TEM, JDB; Writing-Review & Editing: all authors; Supervision: JDB; Funding Acquisition: CALS, JDB.

